# Heterologous Prime-Boost with Immunologically Orthogonal Protein Nanoparticles for Peptide Immunofocusing

**DOI:** 10.1101/2024.02.24.581861

**Authors:** Sonia Bhattacharya, Matthew C. Jenkins, Parisa Keshavarz-Joud, Alisyn Retos Bourque, Keiyana White, Amina M. Alvarez Barkane, Anton V. Bryksin, Carolina Hernandez, Mykhailo Kopylov, M.G. Finn

## Abstract

Protein nanoparticles are effective platforms for antigen presentation and targeting effector immune cells in vaccine development. Encapsulins are a class of protein-based microbial nanocompartments that self-assemble into icosahedral structures with external diameters ranging from 24 to 42 nm. Encapsulins from *Mxyococcus xanthus* were designed to package bacterial RNA when produced in *E. coli* and were shown to have immunogenic and self-adjuvanting properties enhanced by this RNA. We genetically incorporated a 20-mer peptide derived from a mutant strain of the SARS-CoV-2 receptor binding domain (RBD) into the encapsulin protomeric coat protein for presentation on the exterior surface of the particle. This immunogen elicited conformationally-relevant humoral responses to the SARS-CoV-2 RBD. Immunological recognition was enhanced when the same peptide was presented in a heterologous prime/boost vaccination strategy using the engineered encapsulin and a previously reported variant of the PP7 virus-like particle, leading to the development of a selective antibody response against a SARS-CoV-2 RBD point mutant. While generating epitope-focused antibody responses is an interplay between inherent vaccine properties and B/T cells, here we demonstrate the use of orthogonal nanoparticles to fine-tune the control of epitope focusing.

**Table of Contents graphic:** 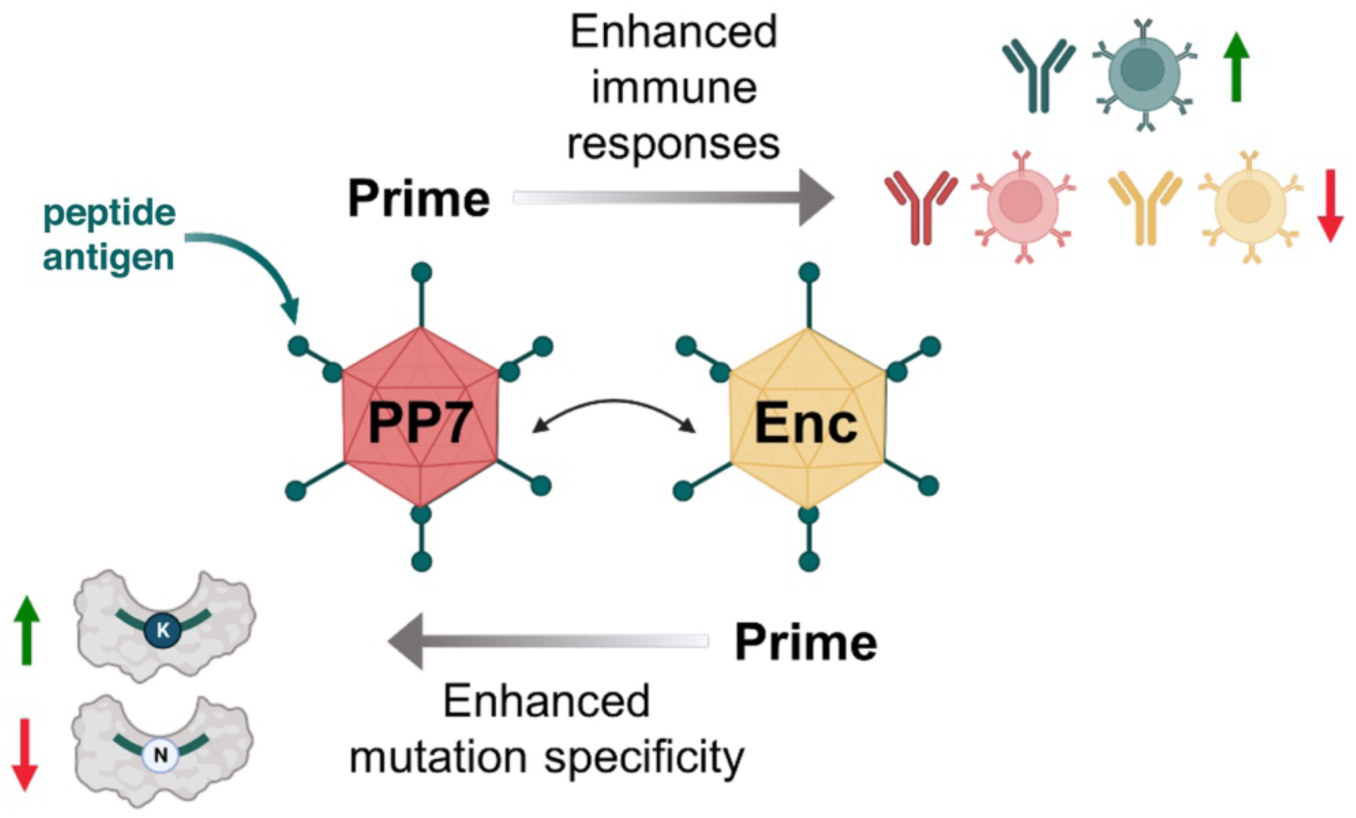

## Introduction

Encapsulins are microbial proteinaceous nanocontainers that sequester functional protein cargos, such as ferritin-like proteins, hemerythrins, and a variety of enzymes within their luminal spaces.^1–7^ Selective protein encapsulation is accomplished by specific protein-protein interactions between peptide sequences present on either the N- or C-terminus of the cargo protein and hydrophobic binding clefts located on the interior faces of the corresponding encapsulin.^4, 8–9^ Since their discovery in the 1990s, encapsulin scaffolds have been derivatized to serve a variety of biotechnology, therapeutic, and materials functions, including as photo-switchable imaging,^10^ catalysis,^11–13^ cellular imaging,^14–16^ targeted drug delivery,^17–18^ toxin remediation,^19^ and vaccination.^20–24^

In modern immunization strategies, the selection of antigenic peptide epitopes from a given pathogen’s protein repertoire is often key to developing successful antibody and T cell responses against infectious pathogens. However, vaccination with such peptides alone usually fails to promote sufficiently robust responses,^25–26^ requiring the use of carrier proteins for conjugate vaccines, a role for which protein nanoparticles are well suited by virtue of their efficient lymphatic trafficking, engagement of immune cell receptors, uptake, processing, and induction of cellular signaling.^25, 27–29^

While encapsulin particles have several beneficial attributes, including high stability, versatile tolerance of sequence modification, and excellent expression yields, they have only occasionally been employed as immunogenic carrier proteins, as follows. A recent report from Care and colleagues showed that a single administration of encapsulin particles from the bacterium *Thermotoga maritima* in BALB/c mice was benign, producing no signs of physical distress or increases in the serum levels of several proinflammatory cytokines.^30^ The same *T. maritima* encapsulins were used to recombinantly display foreign protein domains such as the Matrix protein 2 ectodomain of influenza A virus^21^ as a protomer loop insertion and a truncated form of the gp350 receptor-binding domain from Epstein-Barr virus as a linear extension from the protomer’s C-terminus.^20^ The former particle was administered to mice with Freund’s adjuvant and the latter with the Sigma adjuvant system. Encapsulins from *Myxococcus xanthus* were recently engineered to display receptor binding domains form SARS-CoV-2 virus^24^ or influenza hemagglutinin,^23^ and were tested immunologically with a squalene-in-water emulsion adjuvant. Lastly, only one report has appeared of an encapsulin-peptide conjugate vaccine, in this case using the prototypical Ova2 peptide (SIINFEKL) antigen attached by chemical means to *T. maritima* encapsulins and administered with poly (I:C) adjuvant.^22^

We sought to assess the immunological properties of encapsulin-based vaccines in the absence of co-administered adjuvants, instead achieving adjuvant effects by engineering the platform to package single-stranded RNA (ssRNA) from the bacterial production host to stimulate endosomal Toll-like receptors (TLRs) 7 and 8.^31–33^ Several recombinantly-generated virus-like particle (VLP) vaccine scaffolds derived from ssRNA phages have exhibited enhanced immunogenic profiles due to the self-adjuvanting properties of their nucleic acid cargos.^34–37^ Recently, Kwon and Giessen reported the successful encapsulation of foreign RNAs within the lumen of several encapsulin species by genetically appending a short cationic peptide tag onto the N-termini of the respective encapsulin protomers.^38^ In the work described here, we achieved RNA packaging within encapsulin nanocontainers derived from the mesophilic soil bacterium *Myxococcus xanthus* following a similar peptide-fusion strategy and assessed the immunogenic profiles of the RNA-packaging variants compared to the wild type encapsulin particles. We also inserted a 20 amino acid peptide derived from the receptor binding domain (RBD) of the SARS-CoV-2 spike protein into an exposed surface loop on both the wild type and engineered encapsulin protomers to generate candidate nanoparticle conjugate vaccines for evaluation.

While protein nanoparticle conjugates confer many beneficial properties to candidate vaccines, repeated immunizations are typically necessary to establish strong humoral responses and to induce immune memory towards fused peptide epitopes. There are at least two serious drawbacks associated with such repeat immunizations: (1) the development of potent immune responses to the nanoparticle vehicle itself, leading to rapid clearance of the vaccine candidate from circulation;^39^ and (2) presentation of immunodominant epitopes by the protein nanoparticle vehicle, which partially or wholly mask the presentation of the intended peptide epitope to immune effector cells.^40–41^ The latter phenomenon has been further elaborated by Victoria *et al*. with the concept of “primary addiction,” based on the observation that the cohort of native B cells engaged and activated by the primary vaccine dictates which B cell clones will be chosen for both expansion and seeding of secondary germinal centers after booster vaccinations.^42^ This poses a challenge to conjugate vaccines when the non-targeted scaffold epitopes are competitively antigenic with the desired epitope. One strategy to avert primary addiction is heterologous prime-boost vaccination, in which the same peptide epitope is presented to immune cells on different protein scaffolds over the course of the immunization schedule.^43–45^ In so doing, the proliferation of B cells directed against the vaccine scaffolds is ideally minimized while the immune response directed against the common peptide epitope is concurrently maximized. Here we attempted to focus immune responses^46^ against a 20-mer SARS-CoV-2 peptide by combining our engineered encapsulin scaffolds with a previously reported variant of the PP7 RNA phage VLP in several heterologous prime-boost regimens.

## Results and Discussion

### Design and characterization of RNA-packaging encapsulin variants

The wild type *M. xanthus* encapsulin (Enc) assembles homogenously *in vivo* from 180 copies of a 31.7 kDa protomer subunit to generate a 32 nm, *T* = 3 symmetry icosahedral protein nanoparticle (PNP) that sequesters intracellular Fe^2+^ ions via several packaged ferritin-like proteins.^7^ To effect RNA encapsulation, we genetically appended a hexahistidine (His_6_) affinity tag onto the N-terminus of the encapsulin protomer, which is luminally-oriented in the assembled nanocontainers (Figures 1A and S1). This variant, referred to as HEnc, formed particles morphologically identical to Enc, composed of both *T* = 3 and smaller *T* = 1 populations (Figure 1B and S2), as has been noted previously for Enc nanoparticles generated in the absence of its native cargo proteins.^7, 47^ Native and denaturing agarose gel electrophoresis showed that the HEnc variant packaged host cell RNA during recombinant protein expression, including the 23S and 16S rRNAs observed previously by Kwon and Giessen (Figures S3, S4).^38^ RNA capture in this variant is presumably achieved via nonspecific electrostatic interactions between the 1,080 internally facing histidine residues in each particle and the anionic RNA molecules. By comparison, the wild type Enc encapsulins packaged little to no RNA (Figure S3).

**Figure 1.**
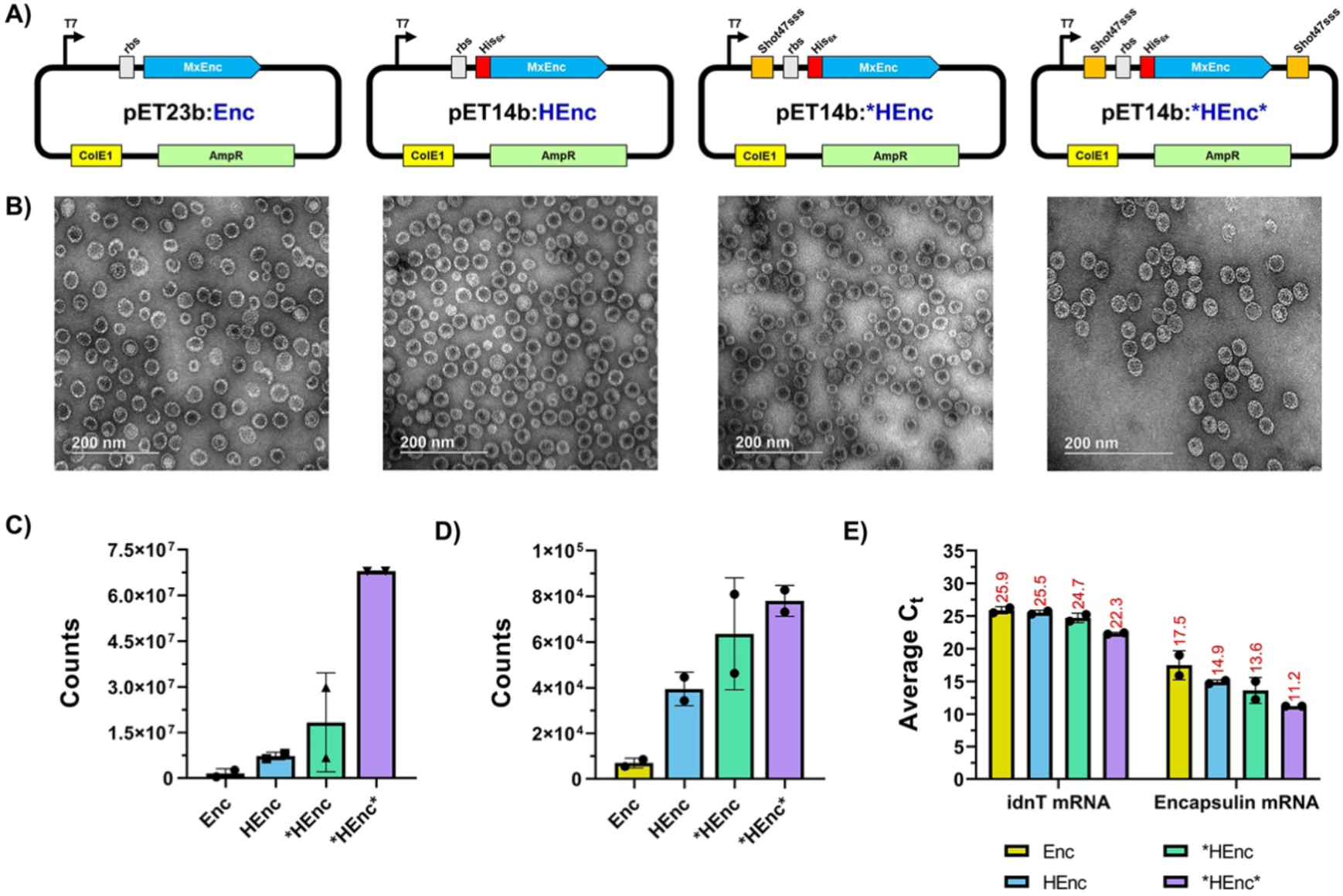
Design and characterization of encapsulin variants. A) Plasmid designs for Enc, HEnc, *HEnc, and *HEnc* variants. (B) Representative TEM images for the indicated particles. C) RT-qPCR assessment of encapsulin protomer mRNA captured within nanoparticles. D) RT-qPCR assessment of *E. coli* idnT mRNA packaged within particles as a proxy for host cell RNA capture during *in vivo* PNP assembly. E) Comparison of the cycle threshold (Ct) values obtained for each sample during RT-qPCR assessments. Data presented in panels C-E represent two independent biological replicates.

We also sought to increase the total amount of encapsulin-packaged RNA by specific, rather than electrostatic, interactions. We have previously biased protein and RNA packaging in VLPs by virtue of paired peptide/anti-peptide aptamer interactions,^48^ and Hilvert and coworkers have encapsulated mRNA within an engineered *Aquifex aeolicus* lumazine synthase (AaLS) variant using a similar method.^49–50^ Here we employed the 37 nucleotide RNA aptamer Shot47sss developed by Tsuji, *et al.*, which binds to polyhistidine affinity tags with picomolar affinity.^51^ Thus, the DNA sequence encoding the anti-polyhistidine aptamer was inserted into either the 3’-untranslated region, or both the 3’- and 5’-untranslated regions of the plasmid encoding our HEnc variant so that mRNA transcribed from the plasmid would possess either one or two copies of the functional RNA aptamer (Figure 1A). Encapsulins expressed from these plasmids (referred to as *HEnc and *HEnc* to denote the use of one or two copies of the Shot47sss aptamer, respectively) also packaged RNAs and were likewise morphologically identical to the Enc and HEnc PNPs, including the production of heterogeneous *T* = 1 and *T* = 3 particle populations (Figures 1B, S2, and S3).

Extraction of RNA from the purified encapsulin particles followed by denaturing gel electrophoresis revealed RNA bands from *HEnc and *HEnc* consistent with the size of *in vitro* transcribed mRNA from the HEnc vector, though a clear band of the same size was not obvious for the HEnc RNA sample (Figure S4). cDNA generated from each extracted RNA using oligo poly-dT primers designed to target the 3’ poly-A tails of cellular mRNA strands was amplified by PCR using primers designed to bind to the 5’ end of the Enc gene and to the T7 terminator originating from the plasmid. Amplicons of the anticipated size were observed for all three RNA-containing encapsulin samples, indicating successful capture of encapsulin protomer mRNA during nanoparticle assembly *in vivo* (Figure S5). Real-time quantitative PCR (RT-qPCR) assessments of the nanoparticle-extracted RNAs using protomer-specific primers confirmed that all of the His_6_-tagged encapsulins packaged their own coding mRNA (Figures 1C, D; standard curves for RT-qPCR quantitation depicted in Figure S6). While the HEnc and *HEnc encapsulins packaged similar amounts of the target mRNA, the *HEnc* PNP containing two copies of the Shot47sss aptamer captured substantially more.

Interestingly, qPCR with primers complementary to the mRNA generated from transcription of the *E. coli* idnT gene (UniProt P39344, used to probe the level of host cell RNA non-specifically incorporated within the engineered encapsulin variants) showed that the level of packaged idnT mRNA followed a very similar trend to the encapsulin mRNA. HEnc, *HEnc, and *HEnc* particles packaged successively greater amounts of mRNA, albeit in quantities 100-fold less than the corresponding encapsulin mRNAs (Figure 1C). Thus, the presence of two Shot47sss aptamers in the protomer mRNA leads to better target mRNA encapsulation, which also results in enhanced sequestration of other cellular RNAs, perhaps through transient hybridization events during nanoparticle assembly. For immunological assessments, we abandoned the *HEnc particle, since the HEnc scaffold packaged very similar amounts of mRNA.

### Immune responses to homologous prime-boost immunizations

Immunization of 7- to 14-week-old female BALB/c and C57BL/6 mice with a total of 3 doses of all encapsulin immunogens was well tolerated (Figure 2A, B). BALB/c mice were given 50 µg of PNPs every 2 weeks and generated significant anti-encapsulin IgG responses after the initial priming with either Enc or HEnc PNPs; total IgG titers continued to rise with each subsequent boost (Figure 2C). While the unmodified Enc PNPs prompted marginally lower IgG responses during the initial weeks of immunization, there was no significant difference between the responses to the two PNPs by week 8. C57BL/6 mice produced higher titers overall and prompted a greater response to the RNA-containing PNP, although this particle was also administered at a higher dose in this preliminary assessment (Figure 2D).

**Figure 2.**
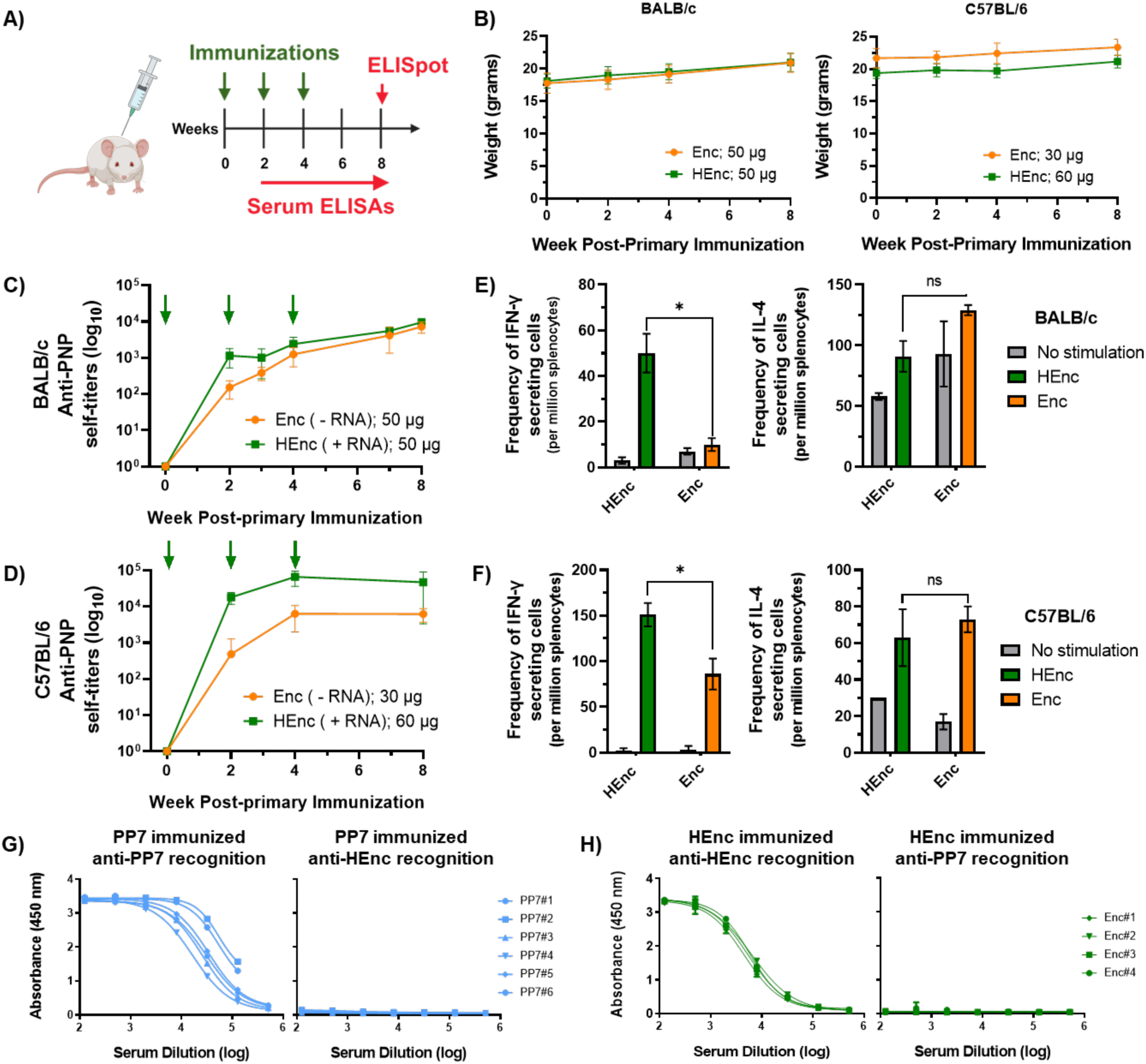
Immunogenicity of unadjuvanted encapsulin particles and cross-recognition with the PP7 scaffold in inbred mouse strains. A) Immunization schedule for native and His-tagged encapsulin variants (Enc and HEnc, respectively). B) Mouse weights throughout immunization regimens. C,D) Self-recognition humoral responses to immunized encapsulins in BALB/c and C57BL/6 mice, respectively; green arrows indicate administered PNP doses. Cellular IFN-γ and IL-4 recall responses at week 8 for BALB/c and C57BL/6 mice are presented in E) and F), respectively. G) Anti-PP7 response and cross-recognition of the HEnc scaffold at week 2 in BALB/c mice immunized with the dimeric PP7 VLP. H) Anti-HEnc response and cross-recognition of the dimeric PP7 scaffold at week 7 in BALB/c mice immunized with the HEnc variant. Each curve represents an individual mouse. All quantitative data in figure displayed as the mean ± SD, and significance analyzed as *p < 0.05 by two tailed t-test.

As a comparison immunogenic platform, we also immunized mice with an engineered version of the virus-like particle derived from the *Leviviridae* phage PP7, generated from a head-to-tail genetic fusion of two copies of the 14.0 kDa coat protein separated by a 4 amino acid AYGG linker^52^ (the dimeric PP7-PP7 particles are designated here as “PP7”). This particle exhibits similar morphological characteristics to the encapsulins (both particles are assembled from multiple coat proteins to form icosahedra of similar size), retains the native ssRNA-packaging ability of the parental PP7 phage, and is conveniently engineered to display foreign peptides within the 4 amino acid junction loop.

Both Enc and HEnc immunizations generated all four major IgG subclasses (IgG1, 2a, 2b/ 2c and G3), characteristic of both T cell dependent and independent pathways. However, PP7 particles proved to be significantly more immunogenic than either encapsulin particle, generating higher titers of self-recognizing total IgG in both BALB/c and C57BL/6 mice (Figure S7A, B). Similarly, the IgG2a/IgG1 ratio, generally an indicator for either Th1 pro-inflammatory (with ratios > 1, our desired outcome for protection against intracellular pathogens^53^) or Th2 anti-inflammatory immune responses (ratios < 1), was significantly higher for PP7 particles compared to both Enc and HEnc in BALB/c (Figure S7C). The same subclass trend was observed in C57BL/6 mice for PP7 VLPs vs. the native Enc PNPs, but not for HEnc particles: these mice responded very similarly to both PP7 and HEnc (Figure S7C). ELISpot analyses of splenocytes collected at week 8 showed significantly stronger antigen-dependent pro-inflammatory IFN-γ production by the HEnc (vs. Enc) particles in both inbred mouse strains, consistent with a greater Th1 contribution^54^ when RNA is packaged (Figures 2E, F).

Heterologous prime-boost immunization with encapsulin and PP7, illustrated in Figure 3B, could be hindered if the two PNP platforms engender cross-reactive immune responses against one another. Fortunately, these carriers were found to be immunologically orthogonal in ELISA assays: no serum antibody cross-reactivity was observed in either direction (i.e., serum from mice immunization with PP7 yielded no detectable recognition of plated Enc, and serum from HEnc-immunized mice did not recognize plated PP7; Figure 2G, H). This result was anticipated given that encapsulins and *Leviviridae* phages originate from evolutionarily distinct protein families and have dissimilar tertiary coat protein structures. In contrast, PP7 and its *Leviviridae* relative Qβ, which share a small but significant amount of amino acid sequence identity (∼20%), have common B cell epitopes that are cross-recognized in serum samples collected from VLP-immunized mice (Figure S8). [We note that a recent report shows no immunological cross-reactivity in BALB/c mice between encapsulins from *T. maritima* and *Quasibacillus thermotolerans*, in spite of the existence of approximately 22% sequence identity between these proteins.^30^ These encapsulin cousins are quite different in size, the former being 24 nm in diameter (*T* = 1) and the latter 42 nm (*T* = 4), although we do not assume that this is the source of their immunological orthogonality.]

**Figure 3.**
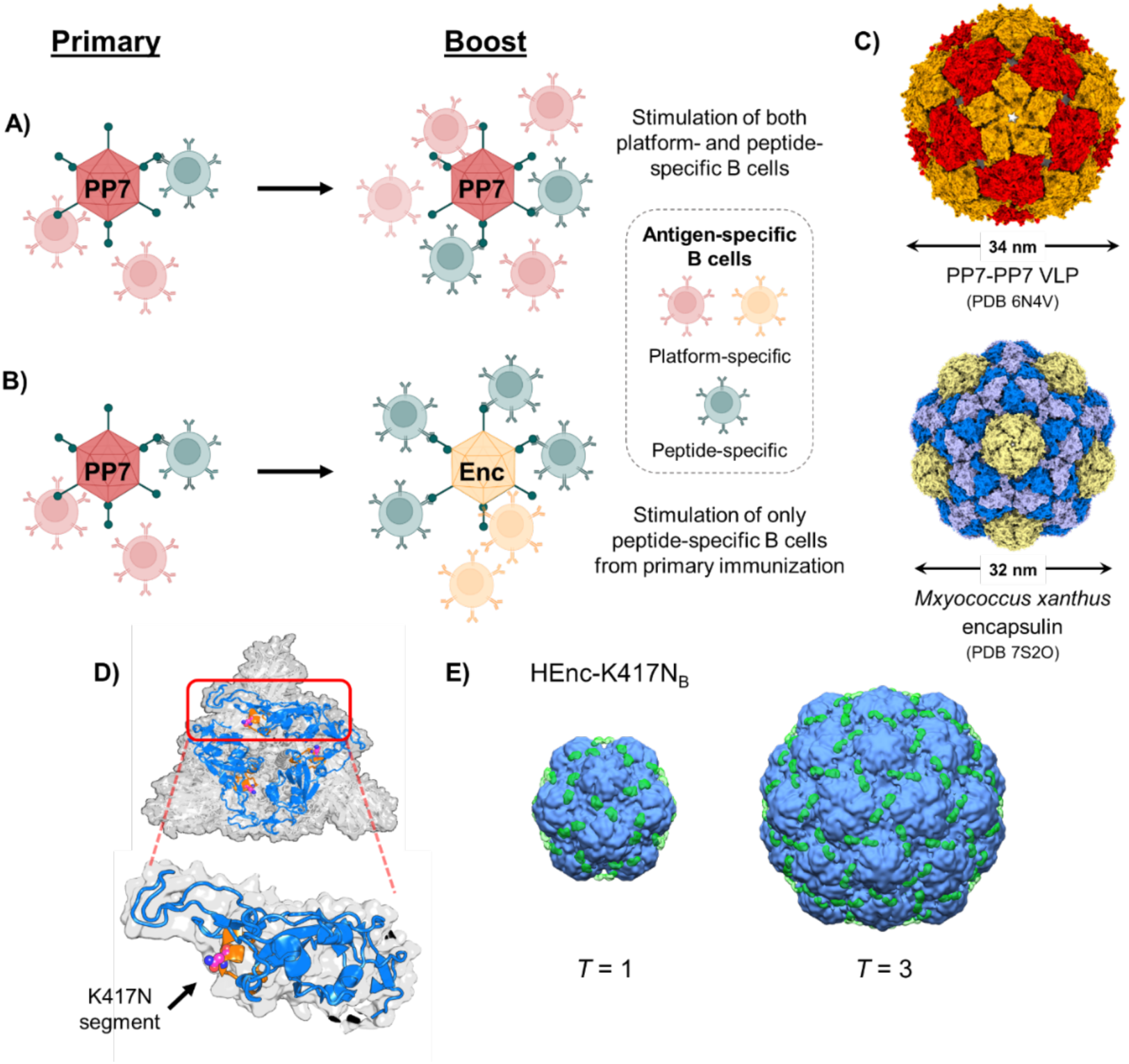
Rationale for alternating protein nanoparticle scaffolds displaying the same antigenic peptide epitope. A) Homologous immunization strategy in which the same nanoparticle scaffold is used in both the primary and booster vaccinations, leading to the proliferation of both peptide and scaffold-specific B cells. B) Heterologous immunization strategy in which the same peptide is present on different nanoparticle scaffolds to selectively expand the population of peptide-specific B cells. C) Atomic structures of the two protein nanoparticles employed herein. D) Structure of the trimeric SARS-CoV-2 spike protein (PDB 8CSA, top-down view) with the individual receptor binding domains depicted in blue and the 20-mer “K417N” peptide sequence depicted in orange. The actual K417N point mutation is depicted with the carbon atoms of the asparagine sidechain represented as magenta spheres. E) Cryo-EM electron volume maps (10Å resolution) of HEnc-K417NB particles depicting the additional electron density (green electron clouds) present on the surfaces of both *T* = 1 and *T* = 3 PNP structures.

### Characterization of encapsulins displaying a foreign SARS-CoV-2 peptide

As the test antigen, we choose a 20-mer peptide sequence (VRQIAPGQTG**<u>N</u>**IADYNYKLP) (Figure 3D) from residues 407 to 426 of the SARS-CoV-2 virus spike protein receptor binding domain (RBD), containing the lysine-to-asparagine mutation at position 417 (i.e., K417N) observed in many of the variants of concern displaying increased infectivity that have emerged since the onset of the COVID-19 pandemic.^55^ In order to mimic native peptide presentation on the progenitor pathogen, we took advantage of the ability of each scaffold to display the peptide polyvalently in a repeating surface loop. For encapsulins, several sites have been identified as potentially viable insertion points for foreign peptides.^18, 21, 56–58^ We tested the insertion of the 20-mer peptide into the encapsulin scaffold at three positions: between residues G58/V59 (designated insertion point A, IP_A_; particle designation Enc-K417N_A_), between residues R135/L136 (IP_B_; Enc-K417N_B_) or between residues N145/G146 (IP_C_; Enc-K417N_C_) (Figure S9A, all residue numbering based on UniProt entry Q1D6H4)). Peptide insertion at IP_A_ and IP_B_ produced intact recombinant particles in good yields, but no such particles were isolated when the peptide was inserted at IP_C_ (Figure S9B). Similarly, a complementary PP7-PP7 particle was generated with the 20-mer peptide inserted between the two glycine residues of the AYGG junction connecting the two coat proteins (designated PP7-K417N-PP7; characterization data in Figure S10).

The IP_A_ and IP_B_ peptide-added encapsulins were also composed of two populations containing *T* = 1 and *T* = 3 particle assemblies (as shown by DLS and TEM, Figures S11 and S12), with no observable differences between the Enc and HEnc PNPs bearing the K417N peptide. The HEnc-K417N_A_ and HEnc-K417N_B_ particles were characterized by cryogenic electron microscopy (cryo-EM): difference maps calculated between the HEnc (*T* = 1 and *T* = 3) and corresponding HEnc-K417N_A_ and HEnc-K417N_B_ structures revealed extra densities for the loop-insertion constructs (Figure S12B and 3E, respectively). These densities closely aligned to the intended IP_A_ and IP_B_ peptide insertion sites and are therefore assigned to the grafted 20mer K417N peptide. Notably, the surface density map for the HEnc-K417N_A_ particles was significantly less defined than the accompanying map generated for HEnc-K417N_B_ PNPs, perhaps due to IP_A_ residing slightly deeper in the clefts surrounding the encapsulin 3-fold symmetry axes.

Subtraction of the cryo-EM Enc map from the HEnc and *HEnc* maps revealed protruding electron densities on the luminal surfaces of the HEnc particles at positions corresponding to the locations of the protomer N-termini (Figure S13). The same N-terminal electron densities were also observed in Enc-subtracted *HEnc* particle maps (not shown), validating the assignment of these volumes to the N-terminally fused His_6_ affinity tags. Interestingly, the Enc, HEnc, and *HEnc* particles all showed several small subpopulations of non-icosahedral particle morphologies corresponding to D5-h15 (150 protomers), D6-h8 (104 protomers), tetrahedral (Th-h4, 84 protomers), D3-h3 (78 protomers), and C2-h9 (114 protomers) symmetry structures in addition to the expected *T* = 3 (180 protomers) and *T* = 1 (60 protomers) particle symmetries (Figure 4A – E). While similar D3 and D5 symmetry arrangements have recently been reported to emerge in several other VLP species following modification of viral capsid proteins,^59–60^ neither these morphologies, nor the other non-icosahedral assembly states observed here have been previously described for either wild type or modified encapsulins. (Recently, distortion of the *M. xanthus* encapsulin shells into undefined, prolate-like cage structures was reported following *in vivo* packaging of PNPs with high levels of proteinaceous cargo, though no explicit symmetry states were refined for the non-icosahedral particles in this work.^61^)

**Figure 4.**
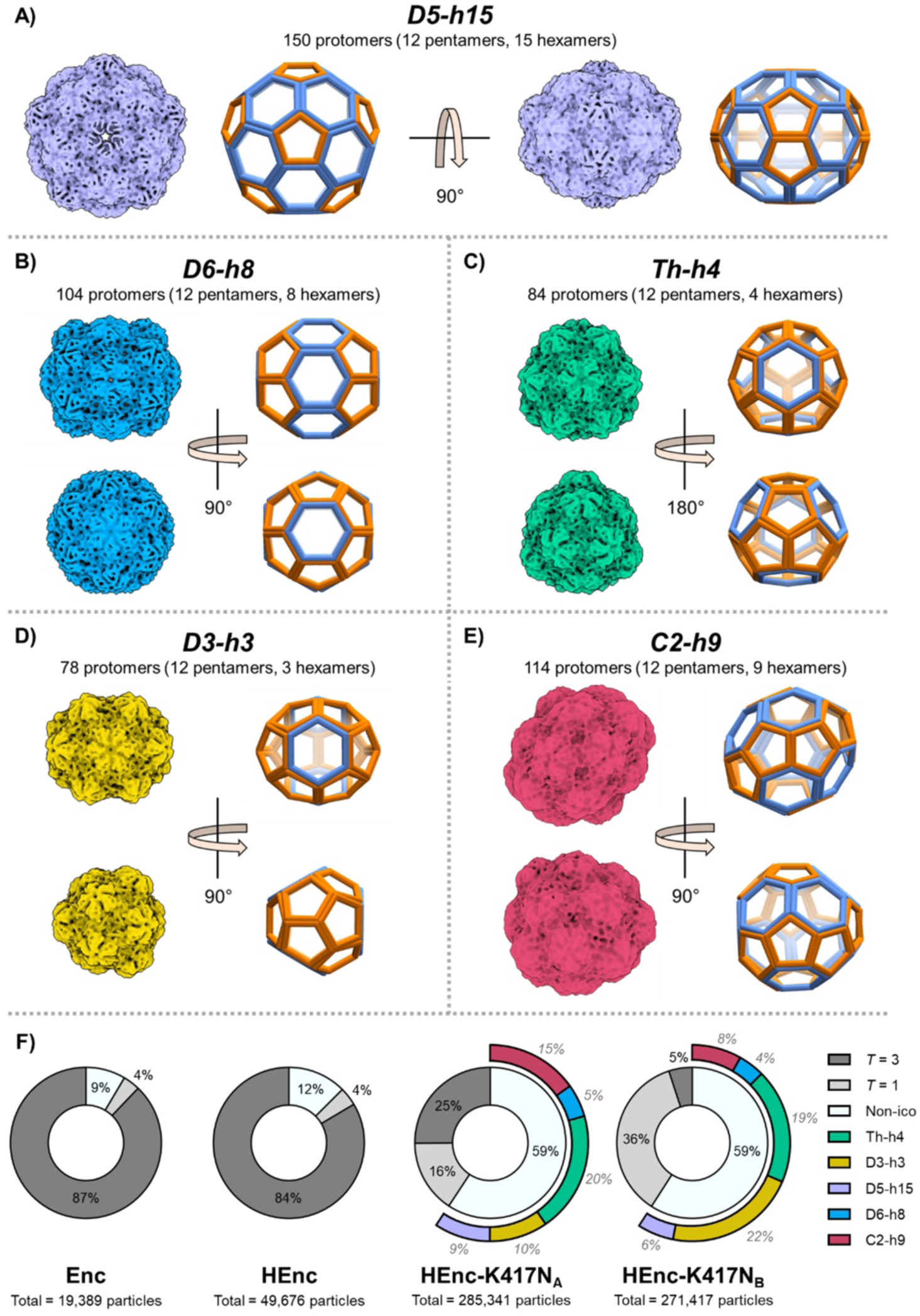
Representative 7-10 Å electron volume maps depicting the observed A) D5-h15, B) D6-h8, C) tetrahedral (Th-h4), D) D3-h3, and E) C2-h9 non-icosahedral cage morphologies in encapsulin PNPs samples (generated from HEnc-K417NA data but found in all samples). All particle morphologies are depicted to scale with one another, and particle names denote symmetries and the number of hexameric tiles observed in each structure (e.g., D5-h15 = D5 symmetry with 15 total hexameric tiles). Wireframe cage models accompany each volume map for clarity with pentameric caps depicted in orange and hexameric tiles depicted in blue. The total number of protomers per cage was determined by counting the number of pentamers and hexamers for each structure and applying the following formula: # of protomers = (# of hexamers*6) + (# of pentamers*5). Distributions of the different icosahedral and non-icosahedral morphologies for each cryo-EM sample are depicted in F). The morphology distributions presented are based on 3D refinements of observed particles following an initial 2D classification analysis. For the HEnc-K417NA and HEnc-K417NB PNPs, the non-icosahedral portions are further divided into subsections representing the populations of each non-icosahedral species. The numerical percentage of each subsection written in lighter, italicized text to distinguish them from the *T* = 3, *T* = 1, and non-icosahedral percentages (black, non-italicized text).

Three-dimensional (3D) classification analyses of cryo-EM data for Enc, HEnc, HEnc-K417N_A_ and HEnc-K417N_B_ PNPs revealed several intriguing phenomena. The observed non-icosahedral morphologies collectively represent minor populations in Enc and HEnc samples (reported generically as “Non-ico” for these samples in Figure 4F due to an inability to accurately quantify the exact proportions of individual non-icosahedral assembly states with the currently employed instrumentation due to their small numbers within the samples) while the *T* = 3 icosahedra represented the major assembly state for each. However, inclusion of the K417N peptide in the encapsulin backbone at either IP_A_ or IP_B_ led to drastic reductions in the proportions of *T* = 3 icosahedra (Figure 4F). Furthermore, the insertion position of the foreign 20-mer peptide appears to affect the distribution of the disparate particle morphologies differently: *T* = 3 icosahedra represented 25% of the total particle population for HEnc-K417N_A_ particles, but only 5% of HEnc-K417N_B_ particles, whereas *T* = 1 particles comprised 16% of HEnc-K417N_A_ and 36% of HEnc-K417N_B_ populations, respectively.

For the non-icosahedral morphologies, the populations of Th-h4, D5-h15, and D6-h8 symmetric assemblies were elevated in both K417N-containing encapsulins relative to Enc and HEnc PNPs, but were roughly equivalent between the two insertion positions. However, the potato-shaped C2-h9 particles were nearly twice as abundant for the HEnc-K417N_A_ variant while the oblong D3-h3 symmetric particles were twice as abundant for HEnc-K417N_B_. Both K417N peptide-containing encapsulins produced greater proportions of smaller morphologies compared to the dominant *T* = 3 assemblies of the Enc and HEnc samples, which was reflected in analytical SEC chromatograms collected for the variant PNPs (Figure S14). Collectively, these data suggest that the inclusion of the K417N sequence disfavors the wild type *T* = 3 quaternary state during particle assembly *in vivo*, perhaps due to the disruption of key interfacial interactions between adjacent protomers or alterations to the kinetic and/or thermodynamic parameters of global particle assembly.

While we were able to resolve the D5-h12, D6-h8, Th-h4, D3-h3, and C2-h9 non-icosahedral structures to an acceptable degree, several additional non-icosahedral samples were also visible in the K417N-encapsulin samples, including D3-h11 (126 protomers), D2-h12 (132 protomers), and D2-h14 (144 protomers) morphologies (Figure S15). However, these assembly states were observed to occur with a significantly lower frequency compared to other non-icosahedral structures, which prevented their full characterization. As such, the population data for these structures was not included in the distribution plots in Figure 4F. Continued cryo-EM analyses are ongoing to try to resolve these additional structures in higher detail.

### Immunological properties of K417N-variant PNPs

In all cases, the morphological mixtures of PNPs obtained for each encapsulin were used for immunological testing. We found no significant difference in the anti-peptide titers generated from immunization with Enc variants possessing the K417N peptide in either IP_A_ or IP_B_ positions (Figure S12F), and so we employed only the HEnc-K417N_B_ particles in homologous and heterologous prime-boost experiments alongside the orthogonal PP7-K417N-PP7 platform. Immunization of BALB/c mice with Enc-peptide particles generated good anti-peptide IgG titers that increased with each booster dose (Figure 5A-C). Though there were no significant differences in anti-peptide (K417N) IgG titer observed following immunization with the peptide-displaying PP7, HEnc, and *HEnc* particles, the non-RNA-containing Enc-K417N particles generated significantly lower anti-peptide responses (Figure 5D). We also tested serum antibodies for cross reactivity towards the full-length RBD protein with K417N mutation, and all groups produced anti-RBD (K417N) responses that closely paralleled the trends observed for the anti-peptide response data, though the titers against the whole protein were 1-2 orders of magnitude lower (Figure 5D). (We note that the molar concentration of RBD (K417N) plated for ELISA (38 nM) was almost 50-fold lower than the biotinylated peptide (2 µM) plated used for anti-peptide analysis, which can affect relative titers.) This significant degree of antibody reactivity with the intact receptor binding domain suggests that the loop conformation of the PNP-displayed peptide resembles that in the polypeptide.

**Figure 5.**
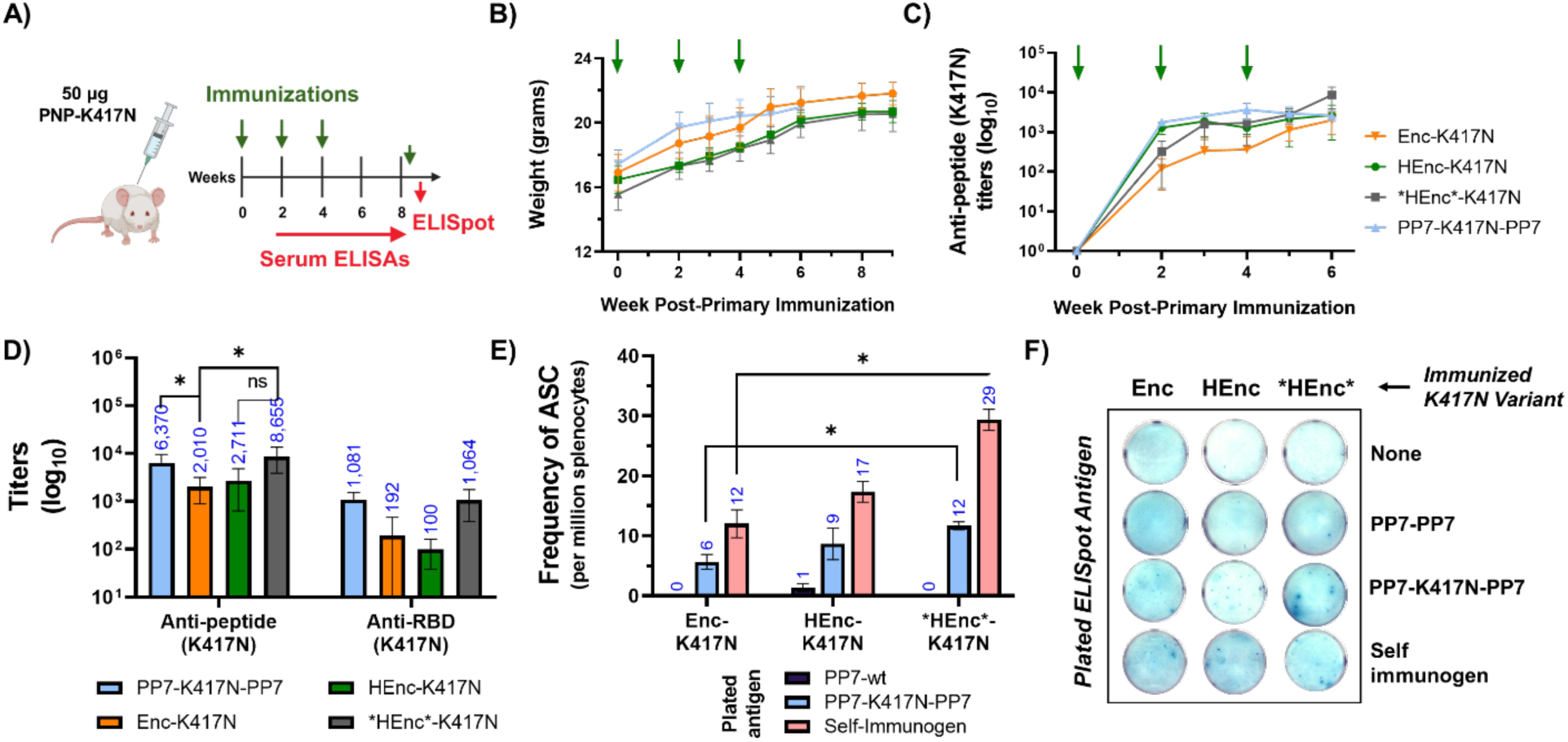
Immunogenicity of Enc-K417NB PNPs A) Immunization schedule in BALB/c mice using either 50 µg of Enc-K417N variants with peptide presented in insertion point B or PP7-K417N-PP7 PNPs. B) Mouse weights throughout the immunization schedule. C) Anti peptide (K417N) titers over time for all immunization groups. D) Anti-peptide and cross-recognizing anti-RBD (K417N) titers at week 6 for mice immunized with either PP7 or Enc variants. E) Frequency of antigen specific, antibody secreting B cells (ASC) from splenocytes analyzed using B cell ELISpot at week 9 after the mice were sacrificed. F) Representative images of the ELISpot wells plated with cells from indicated immunizations atop the indicated plated antigens. N=4 for each group. All quantitative data shows mean ± SEM and significance analyzed as *=p<0.05 by two tailed t-test.

The serum anti-peptide responses for all three Enc-K417N variant immunization groups persisted for at least 15 weeks after the primary immunization with no significant endpoint differences between variants (Figure S16). ELISpot analysis (after stimulation with a final 25 µg dose of the respective K417N-presenting Enc variant three days before fusion) further confirmed that splenocytes extracted from the encapsulin-immunized mice exhibited functional recognition of K417N peptides present in PP7-K417N-PP7 particles, but not the PP7 scaffold alone (Figure 5E, F). Additionally, the frequency of antibody secreting cells recognizing the K417N peptide (presented on PP7 scaffold or as self-immunogen) were significantly lower in Enc-K417N immunized mice compared to mice immunized with *HEnc*-K417N particles (Figure 5E) in good correspondence with serum IgG titer data. As with serum antibody responses, there were no significant differences in the frequency of peptide specific B cells in splenocytes from HEnc and *HEnc* groups.

### Alternate platform heterologous prime-boost immunization

Taking advantage of the immunological orthogonality of the PP7 and encapsulin platforms, we immunized two strains of mice (inbred BALB/c and outbred CD-1 stocks) with either PP7-K417N-PP7 or HEnc-K417N_B_ nanoparticles (abbreviated as “PP7” or “HEnc” for simplicity in this section). The mice were then boosted with either the same or the opposing particle at 2, 4, and 7 weeks after the primary immunization (Figures 6A and 7A). Each heterologous prime-boost sequence was assigned a number (1 to 4, Figures 6B and 7B).

**Figure 6.**
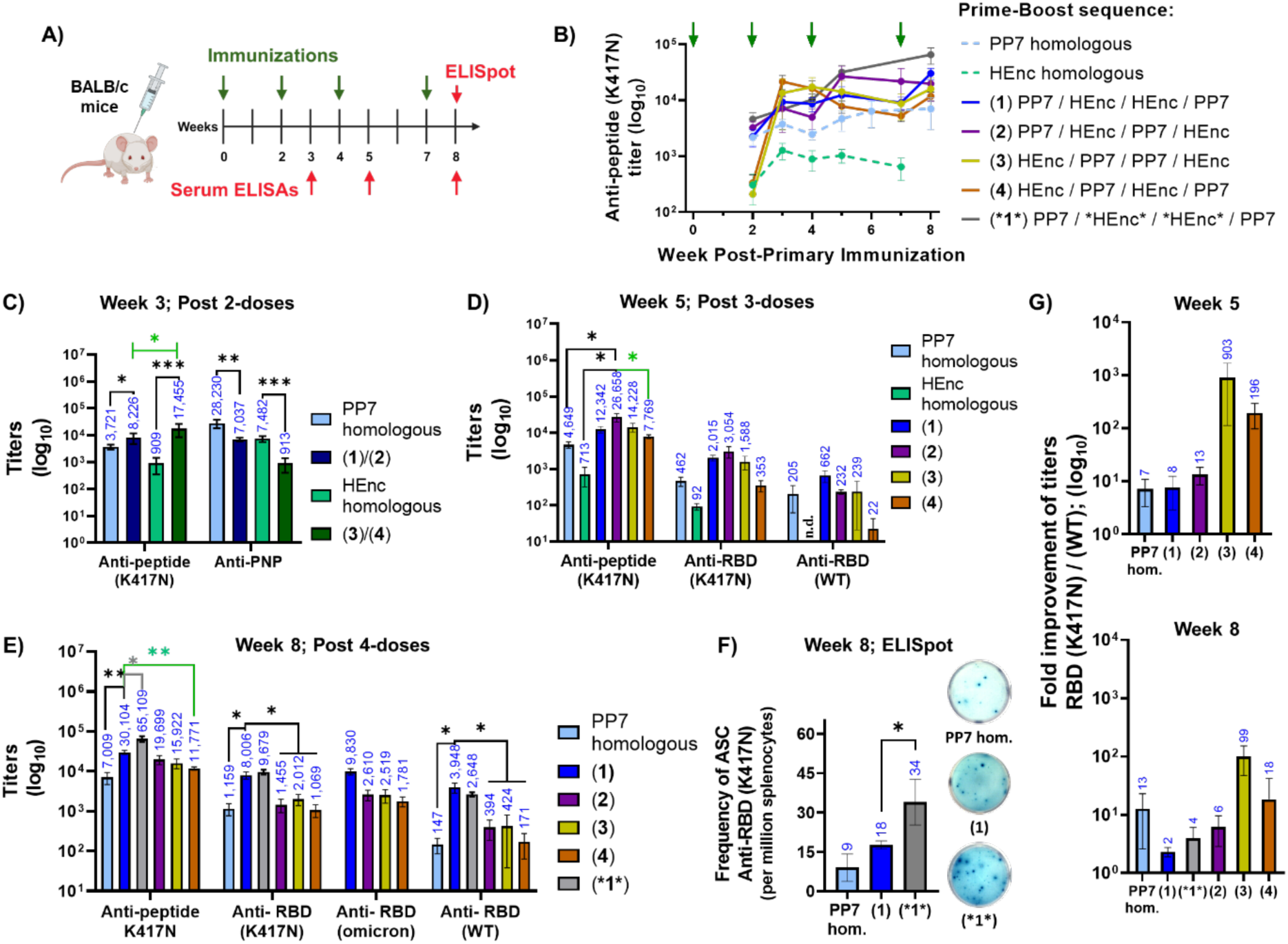
Alternate platform heterologous prime/boost influences antigen processing in BALB/c mice. Immunization schedule with four doses of PNP vaccines displaying K417N peptide, denoted as PP7 and HEnc. Average anti-peptide (K417N) titers per group over time; green arrows indicate successive vaccine doses. Legend indicates numerical codes for heterologous strategies, which are used throughout the rest of the figure. C) Anti-K417N peptide and anti-PNP titers after two doses at week 3. Anti-PP7 responses for PP7 homologous and (1)/(2) groups ; anti-HEnc responses for HEnc homologous and (3)/(4) groups. D) Anti-peptide and cross-recognizing anti-RBD(K417N) titers, anti-RBD(WT) at week 5 after three doses, and E) at week 8 after four doses. F) Frequency of RBD(K417N)-specific antibody secreting splenocytes, at week 8, obtained by using the PP7 homologous, and HEnc-K417N vs. *HEnc*-K417N particles in immunization strategy “1”. Representative ELISpot images from each group are also shown. G) Fold improvement of the titers recognizing RBD(K417N) over RBD(WT) showing immunofocusing at both week 5 and week 8. n.d.= not detectable. N=4 for each group. All quantitative data shows mean ± SEM and significance analyzed as *=p<0.05, **=p<0.005 or ***=p<0.0005 by two tailed t-test.

**Figure 7.**
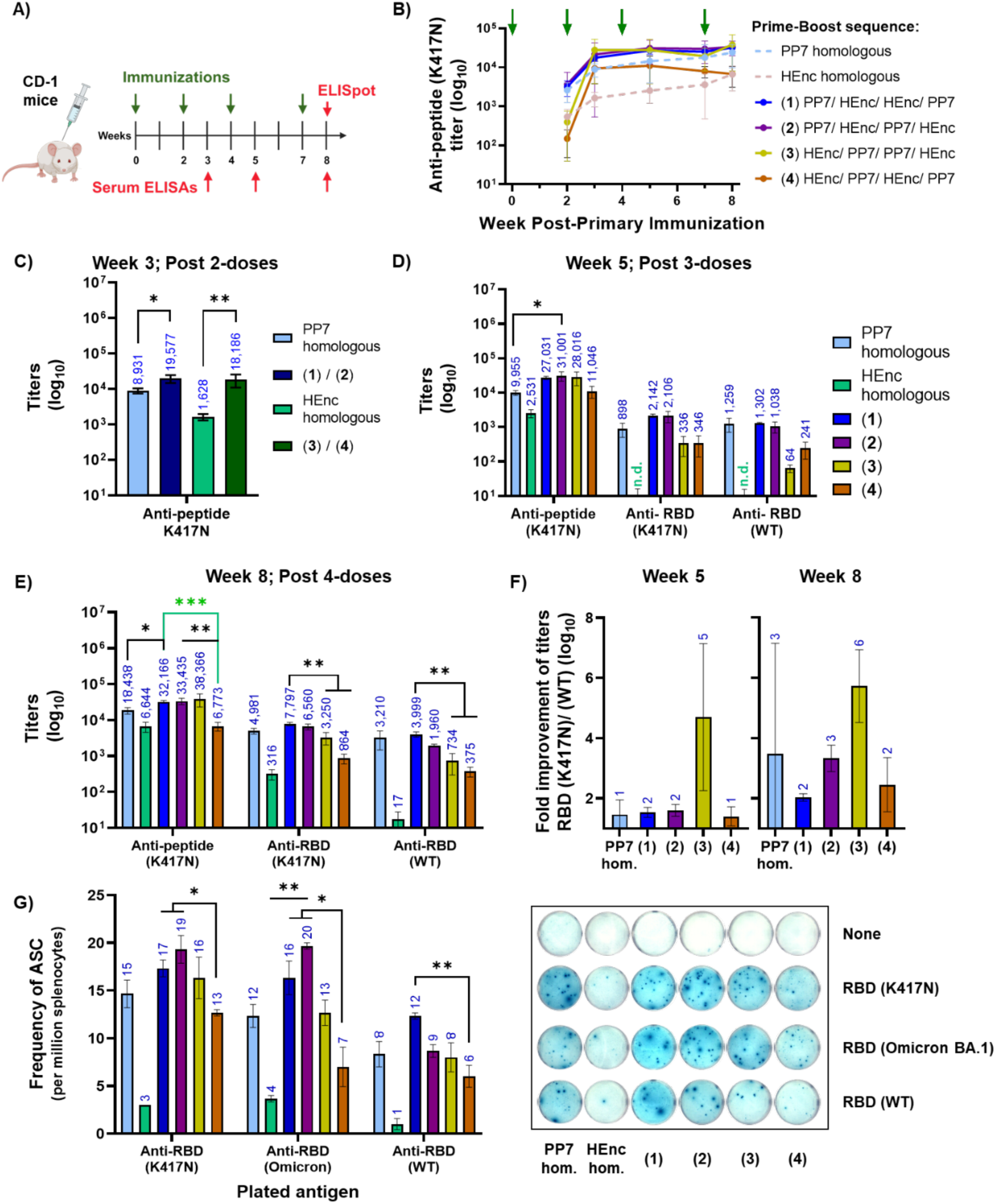
Alternate platform heterologous prime/boost influence antigen processing in CD-1 outbred stocks A) Immunization schedule with total 4 doses of PNP vaccines displaying loop version of K417N peptide antigen denoted as: PP7 and HEnc B) Anti-peptide K417N titers over time; green arrows indicates mice receiving vaccine doses. C) Anti-Peptide (K417N) titers after two doses at week 3. D) Anti peptide and cross-recognizing anti-RBD (mutant), RBD (WT) titers at week 5 after three doses and E) at week 8 after four doses. F) Fold improvement of the titers recognizing RBD (K417N) over RBD (WT) showing immunofocusing at both week 5 and week 8. E) Frequency of RBD(K417N), RBD (Omicron BA.1) and RBD (WT) recognizing antibody secreting cells (ASC) in splenocytes isolated from immunized mice. Representative ELISpot image from each group. n.d.= not detectable. N=4 for each group. All quantitative data shows mean ± SEM and significance analyzed as * = p<0.05, ** = p<0.005 or *** = p<0.0005 by two tailed t-test.

After two doses (prime + one boost), heterologous vaccinations generated significantly higher anti-peptide IgG titers than the corresponding homologous vaccinations in both BALB/c and CD-1 groups (Figures 6C and 7C). This enhancement effect for heterologous strategies was independent of whether the groups were primed to the K417N peptide on the PP7 or HEnc scaffolds. In BALB/c mice, however, HEnc priming led to significantly higher IgG titers than PP7 priming (significance marked in green in Figure 6C). We also observed in BALB/c mice that the heterologous immunizations gave significantly lower anti-PNP responses compared to homologous strategies, evident in serum ELISAs (Figure 6C) and biolayer interferometry (BLI, Figure S17). BLI kinetic responses assessed by immobilizing serum antibodies on anti-mouse capture sensors (AMC) vs. PNPs in solution showed that homologous immunizations with two doses generated significantly lower magnitude responses and lower on-rates (*k*_a_) of serum antibodies with both HEnc and PP7 particles compared to heterologous immunizations. The corresponding off-rates were too slow to assess by this technique.

Continued dosing (second boost = Figures 6D, 7D, and third boost = Figures 6E, 7E) provided higher anti-peptide titers for three of the four heterologous compared to homologous groups. Interestingly, heterologous regimens that began and ended with the more immunogenic PP7 platform – PP7-HEnc-PP7 (approach “2” at week 5) and PP7-HEnc-HEnc-PP7 (approach “1” at week 8) – produced significantly higher anti-peptide titers compared to homologous groups in both BALB/c and CD-1 mice (Figure 6D,E and Figure 7D,E). One heterologous approach (“4” = prime with HEnc and alternate platforms throughout) produced significantly lower anti-peptide (K417N) titers even at week 8 compared to other heterologous schedules and homologous PP7 immunizations in both mouse strains (Figure 6E and Figure 7E). An independent flow experiment using three-doses of either homologous or heterologous prime-boost with three weeks interval between boosts was performed in BALB/c to assess frequency of K417N-peptide specific B cells in lymph nodes and spleens (Figure S21A-D). We observed higher frequency of K417N-peptide reactive FAS/CD95+ GC-derived and (to a lesser extent) CD19+ total B cells after heterologous immunization with PP7 and HEnc-PP7 (approach “2”) compared to PP7 homologous immunization in both lymph node and spleen, although the trends did not reach statistical significance due to the small number of animals used in this preliminary assessment (Figure S21E, F and S22A, B).

The ratio of subclass specific IgG_2a_ and IgG_1_ anti-peptide titers suggested a more balanced Th1/Th2 response for all heterologous groups in BALB/c. (More variation, with strong inclinations towards Th1-type responses were observed in CD-1 mice, Figure S18.) Use of the double-aptamer encapsulin particle (*HEnc*-K417N_B_) in place of the HEnc particles in a heterologous schedule (labeled *1*) gave significantly better anti-peptide titers in BALB/c when compared to both PP7 homologous and heterologous approach “1” (week 5; Figure S17B and week 8; Figure 6E). This is consistent with the better performance of *HEnc* relative to other encapsulin designs in homologous immunization (Figure 5E).

We tested comparative serum binding to three variations of the SARS-CoV-2 receptor binding domain: 1) the RBD from the original Wuhan/HU-1 “wild type” strain [designated RBD(WT)], 2) the RBD containing the K417N point mutation [RBD(K417N)], and 3) the RBD derived from the Omicron (BA.1) variant of concern, containing a total of 15 RBD-localized mutations relative to RBD(WT), including K417N [RBD(omicron)]. In BALB/c mice, the heterologous approach that included PP7 as the initial prime and final boosts, PP7-HEnc-HEnc-PP7 (approach “1” at week 8), produced significantly higher anti-RBD(K417N) and anti-RBD(WT) titers compared to homologous PP7 and the other heterologous approaches (Figure 6E). In CD-1 mice, the trend was similar, although the difference in titer between approaches “1” and “2” did not achieve statistical significance (Figure 7E). Serum antibodies from all heterologous groups maintained similar titers against RBD from the Omicron BA.1 variant in BALB/c mice (not tested in CD-1), despite the additional 14 mutations in the domain (but no additional mutations in the 407-426 epitope presented on the PNP immunogens). Use of the *HEnc* particle in heterologous approach “*1*” provided no further improvement in cross-recognizing anti-RBD serum titers compared to HEnc in the same strategy, but induced a significantly greater frequency of RBD(K417N)-specific B cells in ELISpot assays (Figure 6F).

In contrast to the trend in overall serum titer, which is enhanced by priming with the PP7 particle, the serum antibody repertoire induced by encapsulin-initiated heterologous approaches “3” and “4” (but not the homologous prime-boost regimen) showed over 100-fold better recognition of the RBD protein containing the K417N point mutation relative to the WT sequence at week 5, which persisted at week 8 (Figure 6G). In other words, a more selective peptide-focused antibody response seems to have been generated in BALB/c mice by a less immunogenic primary immunization. This outcome was recapitulated, albeit to a much smaller degree, by week 8 in CD-1 mice (Figure 7F).

The ability of homologous and heterologous immunization to elicit the production of RBD cross-recognizing B cells was assessed by ELISpot in the more diverse outbred CD-1 mice (Figure 7G). Such cells were almost completely absent after homologous HEnc immunization, but fairly robust with all other regimens, including homologous PP7 treatment. Platform-switching after initial PP7 prime immunization (approaches “1” and “2”) was consistently effective, statistically outperforming the alternating HEnc-initiated regimen “4”, but not the regimen in which two PP7 doses followed the HEnc prime (approach “3”). In all cases other than homologous HEnc dosing, the frequency of K417N RBD-responsive B cells was approximately double that of WT RBD-responsive cells (Figure 7G and S20).

## Conclusions

Previously reported immunological studies^20–24^ using encapsulin scaffolds have been augmented here by the encapsulation of immunostimulatory RNA. They can be expected to share the same advantageous properties as virus-like particles for use as vaccine platforms: naturally efficient trafficking to draining lymph nodes due to their size,^27^ polyvalency-enhanced ability to bind native B cells present in circulation,^62^ and ready uptake by antigen presenting cells enhanced by interactions with pathogen-associated molecular pattern receptors.^63^ While less immunogenic than *Leviviridae*-derived VLPs such as PP7, presumably because of the nature of their peptide epitopes, we show here that encapsulins can be advantageous as an immunologically orthogonal carrier protein.

Heterologous prime boost strategies have been used in both pre-clinical and clinical settings. Examples include different transfection vectors and mRNA vaccines expressing the same SARS-CoV-2 antigens^64–65^ and DNA vaccines employed alongside recombinant proteins targeting key epitopes of HIV.^66–67^ Similarly, a DNA vaccine was paired with a modified vaccinia Ankara (MVA) viral vector vaccine to target the malaria strain *Plasmodium falciparum*,^68^ and a different DNA vaccine was used with a tuberculosis-related antigen (a recombinant Bacille Calmette-Guérin protein) to target the *Mycobacterium bovis* strain or tuberculosis.^69^ Another study has shown enhanced B cell recognition of a cross-conserved site in HIV-1 trimer envelope protein (Env) following heterologous immunization with Env proteins with epitopes shielded vs. unshielded with native N-glycans.^70^ Similar beneficial effects of heterologous prime boost on antigen-specific CD4 and CD8 T cells have been widely reported for vaccination targets against HIV^71–72^, malaria,^73–74^ and tuberculosis.^75–76^

To the best of our knowledge, there have been only two prior uses of different protein nanoparticles to focus immune responses towards specific epitopes. The first was an interesting study in which three plant viruses – cowpea mosaic virus, cowpea chlorotic mottle virus, and Sesbania mosaic virus – were all engineered to display a tumor-associated antigen peptide to target HER2+ breast cancers.^45^ This route produced significant reductions in tumor volume and increased survival with a heterologous prime-boost regimen compared to homologous immunization. However, platform cross-reactivity was not investigated, and only a single variation of multiple possible prime-boost formulations was described. The second example targeted the FP8 peptide of the HIV envelop protein by displaying the same peptide on *Leviviridae*-derived Qβ and MS2 VLPs. Similar to our findings, the authors observed significantly higher anti-FP8 peptide-specific titers with the heterologous strategy (Qβ prime, MS2 boost) compared to homologous immunizations with either VLP separately.^43^

The present study introduces or reinforces several key lessons. First, bacterial RNA encapsidation successfully enhanced the immunogenic properties of the encapsulin particle, evident in significantly lower anti-PNP and anti-K417N titers raised by native Enc particles compared to the best RNA-sequestering *HEnc* particles tested (serum and B cell frequency data in Figure 5D and 5E, respectively). However, the encapsulin platform repeatedly generated lower antibody titers when compared to the dimeric PP7-based formulations (HEnc and PP7 homologous groups plotted in Figure 6C and 6D, respectively), and IgG titers tended to decline if HEnc particles were used for booster doses (particularly evident in approaches “2” and “4” in Figure 6B). Similarly, vaccination groups primed with encapsulin particles and boosted with PP7 particles yielded only transiently high IgG titers that then decreased steadily until the subsequent boost (approaches “3” and “4” in Figures 6B and 7B). Finally, pairing the PP7 particles with the more antigenic *HEnc* particles in the most successful PP7-*HEnc*-*HEnc*-PP7 regimen yielded both the strongest and most resilient antibody titers of all the immunization designs tested in BALB/c mice, significantly outperforming the homologous PP7 prime-boost regimen at all major timepoints (Figure 6E, F, and Figure S19 A, B). However, priming with the encapsulin-peptide particle followed by boosting with the PP7 particle showed greater recognition of the K417N point mutation in the context of the full-length RBD domain (Figure 6G). Serum recognition of the peptide sequence in the more relevant BA.1 Omicron variant of the SARS-CoV-2 RBD, which contains 14 additional mutations in addition to the K417N point mutation, was not compromised relative to the variant RBD containing only the K417N mutation (Figures 6E and 7G).

Our observation of favorable immune outcomes using a heterologous platform prime-boost strategy corresponds to expectations based on the Original Antigenic Sin (OAS) theory popularized for influenza virus.^77^ By increasing the antigenic distance between the protein nanoparticles used for primary and secondary immunizations, we selectively encourage development of B cells recognizing target peptide epitopes that were developed after priming. This effect was evident in both increased anti-peptide serum response (Figures 6C and 7C) and increased frequency of splenic B cells recognizing recombinant RBD protein in BALB/c mice in the heterologous groups (Figure 6F), along with increased counts of K417N-peptide reactive GC-B-cells in spleen and draining lymph nodes of mice immunized with heterologous approach “2” (Figure S21E). Additionally, we observed decreases in PNP scaffold-specific serum responses in the initial periods (Figure 6C and S17). This is also supported by a recent report showing that eliciting high titers of high-affinity antigen specific antibodies after primary immunizations leads to limited participation of naive cognate B cells in germinal centers after secondary immunizations.^78^ Among the multiple heterologous prime-boost strategies we tried, it was notable that priming with the less immunogenic encapsulin scaffold resulted in a more specific, mutant-focused anti-peptide immune response (Figure 6G and 7F). This is consistent with the OAS-based assumption that the B cell repertoire produced by the priming immunization sets the course for subsequent amplification and somatic mutation, if priming with the HEnc-peptide particle produces a higher proportion of peptide-focused B cells to be carried into affinity maturation amidst the lower overall response.

## Materials and Methods

### Expression and Purification of Encapsulins

All Enc nanocontainers were expressed and assembled in BL21(DE3) *E. coli* host cells (New England Biolabs) following an identical expression protocol. Briefly, a single transformant bacterial colony was inoculated into 50 -250 mL of 2YT media supplemented with a final concentration of 0.1 mg/mL carbenicillin in a baffled Erlenmeyer flask. The culture flask was subsequently incubated in a 37 °C shaking incubator set at 250 rpm for 18-20 hours without IPTG induction. Cells were then harvested by centrifugation in a JA-16.250 rotor (Beckman Coulter) at 8,000 rpm for 10 minutes and were either processed immediately or stored at -80 °C for future purification.

All encapsulin purifications followed an identical protocol. Cell pellets from the 250 mL cell culture were resuspended in 50 mL of 50 mM Tris-HCl (pH 8.5) buffer supplemented with DNase I and RNase A (Millipore Sigma, final concentration of 2 µg/mL for each). Resuspended cells were lysed by 50 W sonication pulses in an ice-water bath for 10 minutes (5 second sonication pulses with 5 seconds rest between pulses for a total of 5 minutes of active sonication time). Insoluble cellular debris was removed by centrifugation in a JA-17 rotor (Beckman Coulter) at 14,000 rpm for 15 minutes at 4 °C. The supernatant was then passed over two HiTrapTM Q-FF anion exchange chromatography columns connected in a head-to-tail fashion (Cytiva; 5 mL column volume per column; columns pre-equilibrated with 10 column volumes of 50 mM Tris-HCl [pH 8.5] buffer). The column flow-through was collected and an additional 10 mL of lysis buffer was passed over the stacked columns and collected in the flow-through fraction. Next, PEG-6000 and NaCl solids were added to the flow-through fraction to final concentrations of 20 mg/mL and 10 mg/mL, respectively. The sample tube was gently agitated at room temperature until all solids were dissolved, and then the sample was incubated at 4 °C for 1 hour to precipitate nanocontainers. Solid precipitates were collected by centrifugation in a JA-17 rotor at 14,000 rpm for 15 minutes at 4 °C, the resulting supernatant was decanted, and the precipitate pellets were resuspended in 5 – 10 mL of 50 mM potassium phosphate (pH 7.5) buffer. Lipids and other cellular debris were removed via organic extraction with an equal volume of a 1:1 chloroform/*n*-butanol solution. The aqueous layer containing encapsulin nanocontainers was resolved by centrifugation in a JA-17 rotor at 14,000 rpm for 10 minutes at 4 °C, and nanocontainers were then further purified by sucrose density ultracentrifugation (10-40% w/v gradients prepared in 50 mM potassium phosphate buffer at pH 7.5) in a SW-32 rotor (Beckman Coulter) spinning at 28,000 rpm for 4 hours at 4 °C. The gradient section containing assembled encapsulins was removed via aspiration and nanocontainers were isolated in a final ultracentrifugation step in a Type 70Ti rotor (Beckman Coulter) spinning at 68,000 rpm for 2 hours at 4 °C. The resulting encapsulin-containing pellets were resuspended in 50 mM potassium phosphate (pH 7.5) buffer, passed through a 0.2 µm polyetherlsulfone syringe filter, and were either stored at 4 °C for immediate use or were frozen at -80 °C for long-term storage.

### Expression and Purification of PP7-PP7 VLPs

Both unmodified and K417N-containing PP7-PP7 VLPs were expressed and assembled in BL21(DE3) cells, and were subsequently purified in an identical fashion. Briefly, a single transformant colony was inoculated into 0.5 L of 2YT medium supplemented with a final concentration of 50 µg/mL of streptomycin in a baffled Erlenmeyer flask. The culture was grown at 37 °C in a shaking incubator (250 rpms) to an OD_600_ value between 0.8 – 1.0. Protein expression was then induced by adding a final concentration of 1 mM isopropyl ß-D-1-thiogalactopyranoside (IPTG). The cell culture was subsequently incubated at 25 °C overnight prior to harvesting the cells via centrifugation. VLPs were subsequently purified from the harvested cells in a similar fashion to the encapsulins with the following exceptions: 1) the cells were resuspended in 50 mM potassium phosphate (pH 7.0) rather than 50 mM Tris-HCl; 2) VLPs were not subjected to anion exchange chromatography; 3) VLPs were precipitated from the clarified cell lysate using a final concentration of 0.265 mg/mL (NH_4_)_2_SO_4_ rather than PEG-6000/NaCl. All other stages of the purification process were identical to those employed for encapsulins.

### Characterization of Purified Nanoparticles

The concentrations of purified PNP samples were determined via Bradford assays using Coomassie Plus Bradford Assay Kits (Pierce). Bovine serum albumin was employed as the protein standard. All samples were prepared in triplicate at several different dilutions to ensure accurate concentration values. Sample homogeneity was assessed via dynamic light scattering (DLS). Protein samples were prepared at a final concentration of 0.125 mg/mL, and DLS assessment was carried out using a Dynapro DLS plate reader (Wyatt Technologies) in a black-walled 384-well plate. Sample assessment was performed using an instrument laser power between 10 – 20% and an attenuation value between 10 – 30%. Isocratic size exclusion FPLC was performed by injecting 100 µL of protein sample into a hand-poured Surperose 6 column. The column was pre-equilibrated in 50 mM HEPES buffer (pH 7.5) prior to sample loading and the same buffer was pumped over the resin bed at a constant flow rate of 0.4 mL/minute for the duration of the sample run. Absorbance data were collected continuously at 260 and 280 nm on a multiwavelength detector. For transmission electron microscopy (TEM) assessment, protein samples were diluted to a final concentration of 0.05 – 0.1 mg/mL in 50 mM potassium phosphate (pH 7.5) buffer. Individual TEM grids were prepared by applying 8 µL of protein sample onto the carbon surface of 300 mesh lacey formvar/carbon TEM grids (Ted Pella Inc.) for 90 seconds. Grids were then gently blotted perpendicularly against a Kimwipe and residual buffer salts were removed by laying the carbon surface of the grids sequentially on top of two 0.5 mL drops of deionized water (10 seconds incubation per water drop). The grids were again blotted perpendicularly against a Kimwipe and negative staining was performed by applying 8 µL of a 2% w/v uranyl acetate solution onto the grid surface for 45 seconds. The grids were blotted against a Kimwipe one final time and then allowed to air dry for 5 minutes at room temperature. TEM imaging was conducted using a Hitachi HT7700 transmission electron microscope operating at an accelerating voltage of 120 kV.

### Cryogenic Electron Microscopy

All samples were prepared on UltrAufoil^®^ holey grids (300 mesh 1.2/1.3). Grids were plasma cleaned for 7 seconds using an oxygen and argon mix in a Solarus II Gatan Plasma system. The grids were plunge-frozen into liquid ethane using a Thermo Fisher Vitrobot Mark IV system with a blotting time of 1 second and waiting time of 10 seconds with 3 µL of sample deposited onto the grid. The concentrations for each sample were as follows: Enc wildtype at 2.5 mg/mL, HEnc at 3.3 mg/mL, *HEnc* at 3.5 mg/mL, HEnc-K417N_A_ at 4.2 mg/mL, and HEnc-K417N_B_ at 4 mg/mL. All grids were imaged on a Thermo Fisher Glacios 200kV microscope (SEMC Glacios^TM^ 2) equipped with a Falcon3 direct electron detector. Movies were collected in linear mode at 1000 ms exposure time and 25 frames (40 ms/frame) with a pixel size of 1.204 Å and a nominal defocus of -1.5 µm. Data acquisition parameters were as follows: Enc wildtype (session m23aug15a), 1,417 movies with a dose of 50.44 e^−^/Å^2^; HEnc (session m23jul18b), 1,820 movies with a dose of 59.69 e^−^/Å^2^; *HEnc* (session m23aug01a), 1,632 movies with a dose of 48.94 e^−^/Å^2^; HEnc-K417N_A_ (session m23aug21f), 2,971 movies with a dose of 63.91 e^−^/Å^2^; HEnc-K417N_B_ (session m23sep01c), 2,459 movies with a dose of 62.94 e^−^/Å^2^. All data acquisition was performed using Leginon.^79^ Collected movies were first processed with UCSF MotionCor2.^80^ Motion-corrected and dose-weighted micrographs were imported into CryoSPARC v4.3.1,^81^ where CTFFIND4 was used to estimate contrast-transfer function.^82^ Blob picker was used for initial particle picking with a particle size set to 300-330 Å. Particle picks were extracted with a box size of 512 pixels for further processing. 2D classification was used to separate particles of different sizes and ab initio reconstruction was used to obtain initial models and determine particle symmetries. Final refinement was performed with the assigned symmetries applied. Five different cages assemblies were identified in HEnc-K417N_A_ and HEnc-K417N_B_ samples, which were all resolved to < 7 Å resolution. These maps were used as initial models for heterogeneous refinement to obtain final values for 3D classification of whole datasets. For difference map generation, final maps were first low-pass filtered to 10 Å resolution. Density subtraction and visualization was performed in UCSF Chimera 1.13.1.^83^ Volume maps of HEncA-K417N_A_ assemblies from this manuscript were deposited in the EMDB and assigned access ID codes as follows: *T* = 3 symmetry (EMD-43036); *T* = 1 symmetry (EMD-43013); Th-h4 symmetry (EMD-43016); C2-h9 symmetry (EMD-43039); D2-h12 symmetry (EMD-43041); D2-h14 symmetry (EMD-43042); D3-h11 symmetry (EMD-43040); D3-h3 symmetry (EMD-43037); D5-h15 symmetry (EMD-43043); D6-h8 symmetry (EMD-43038). A volume map for the *T* = 3 assembly state of the wild type encapsulin was also deposited (EMD-43113) for reference against previously obtained encapsulin maps.

### RNA Extraction from Encapsulin Samples

Encapsulated RNAs were extracted from purified encapsulins by mixing 200 µL of protein sample with 600 µL of TRIzol^TM^ extraction reagent (Invitrogen). The samples were inverted several times to ensure thorough mixing followed by a 5-minute incubation at room temperature. Next, 120 µL of chloroform was added to each sample and the sample tubes were vortexed briefly before incubating for another 5 minutes at room temperature. The resulting organic and aqueous layers were resolved via centrifugation at 12,000 x *g* for 15 minutes at 4 °C, and the upper aqueous layer from each sample was carefully transferred to a fresh sample tube with a pipette. An equal volume of 100% ethanol was added to each aqueous phase and the samples were mixed gently by inversion. Extracted RNAs were then further purified using Monarch® RNA Cleanup Kits (New England Biolabs) following the manufacturer’s instructions. Captured RNAs were eluted from the Monarch kit affinity columns in 20 µL of DEPC-treated water and were stored at -80 °C until needed. The final concentrations and purities (i.e., A260/A280 and A260/A230 ratios) of isolated RNAs were determined Nano-drop analysis.

### Denaturing Agarose Gel Electrophoresis Assessment of Extracted RNAs

RNA ladders were purchased from New England Biolabs. Extracted RNAs were qualitatively assessed in 2% (w/v) agarose gels prepared in 0.5x TAE buffer containing a final concentration of 0.5 µg/mL ethidium bromide. RNA samples (1 µg each) were mixed with 2x RNA Loading Dye (New England Biolabs), denatured at 72 °C for 10 minutes, and then cooled on ice for 1 minute prior to being loaded into the agarose gel. Electrophoresis was carried out in 0.5x TAE at a constant potential of 120 V for 50 minutes. In vitro transcribed (IVT) mRNA used for comparison in NAGE gels was produced using a HiScribe T7 High Yield RNA Synthesis Kit (New England Biolabs) following the manufacturer’s instructions. Briefly, DNA primers designed to bind to the parent pET14b vector 15 base pairs upstream from the T7 promoter and directly downstream from the T7 terminator sequence, respectively, were used to generate a PCR amplicon to serve as a template for IVT RNA synthesis. The PCR amplicon was purified by sequential gel extraction (1.5% agarose gel prepared in 0.5x TAE) and affinity column chromatography (Monarch® PCR & DNA Cleanup Kit, New England Biolabs). For IVT synthesis, 1 µg of the purified amplicon template was mixed with 3 µL of 10x T7 Reaction Buffer, 3 µL of each ribonucleotide triphosphate stock solution (100 mM stock concentration each ribonucleotide), 3 µL of T7 RNA Polymerase Mix, and 2 µL of DEPC-treated water for a final volume of 30 µL. The IVT reaction was carried out at 37 °C for 2 hours in a thermocycler block followed by removal of the template DNA by adding 60 µL of DEPC-treated water, 10 µL of 10x DNase I Reaction Buffer, and 2 µL of 2000 U/mL DNase I to the IVT reaction mixture (New England Biolabs). Template DNA digestion was carried out at 37 °C for 30 minutes in a thermocycler block prior to purification of the RNA using a Monarch RNA Cleanup Kit (New England Biolabs). Purified mRNA was eluted in DEPC-treated water and stored at -80 °C prior to use. Densitometry analyses of NAGE gels were performed using the FIJI image analysis software package.^84^

### Generation and PCR Assessment of cDNAs

Prior to cDNA synthesis, isolated RNA samples were treated with RNase-free DNase I (New England Biolabs) to remove any contaminant DNAs which might yield false positives in downstream PCR tests. Briefly, 1 µg of encapsulin-extracted RNA was mixed with 1 µL of 10x DNase I Reaction Buffer and then the mixture was brought to a volume of 9 µL using DEPC-treated water. Next, 1 µL of 2000 U/mL DNase I (New England Biolabs) was added to bring the final reaction volume to 10 µL. The reactions were thoroughly mixed and then incubated at 37 °C in a thermocycler block for 30 minutes. The DNase I was subsequently inactivated by ramping the block temperature to 75 °C for 10 minutes.

cDNAs were synthesized from isolated RNA samples using ProtoScript II First Strand cDNA Synthesis Kits (New England Biolabs) following the manufacturer’s instructions. Briefly, 4 µL of DNase I-treated RNA solution (100 ng/µL) were mixed with 10 µL of 2x ProtoScript II Reaction Mixture solution, 2 µL of oligo d(T)23-VN reverse primer, 2 µL of ProtoScript II Enzyme Mixture solution, and 2 µL of DEPC-treated water to reach a final volume of 20 µL. Negative control reactions were prepared in parallel in which the ProtoScript II Enzyme solution was replaced with 2 µL of DEPC-treated water. All reaction mixtures were thoroughly homogenized and then incubated at 42 °C for 1 hour in a thermocycler block. Reverse transcriptase from the enzyme solution was subsequently inactivated by ramping the block temperature to 80 °C for 5 minutes. cDNA samples were stored at -20 °C until needed.

Successful cDNA synthesis was assessed by PCR using a gene-specific forward primer (5’ – GTGCATATGCCGGATTTCCTCGGTC – 3’) and a plasmid-specific reverse primer (5’ – CGTTTAGAGGCCCCAAGGGGTTATGCTAG – 3’). Briefly, 1 µL of sample from an individual cDNA reaction was mixed with 12.5 µL of 2x Q5® High-Fidelity Master Mix (New England Biolabs), 0.5 µL of each primer (0.2 µM final concentration of each primer), and 10.5 µL of sterile water for a final volume of 25 µL. Positive control reactions were prepared in parallel containing 1 µL of 10 ng/µL pET14b:HEnc plasmid DNA in place of the cDNA reaction mixture. PCRs consisted of (1) an initial denaturation step at 98 °C for 2 minutes; (2) 35 cycles of strand denaturation at 98 °C for 15 seconds, primer annealing at 58 °C for 30 seconds, and new strand extension at 72 °C for 30 seconds; (3) a final polishing step at 72 °C for 3 minutes. Following the completion of the PCRs, 10 µL samples were removed from each reaction mixture and loaded into separate wells of a 2% w/v agarose gel containing a final concentration of 0.1 µg/mL ethidium bromide (gel prepared in 0.5x TAE buffer). Electrophoresis was conducted at a constant potential of 120 V for 45 minutes in 0.5x TAE buffer prior to imaging the gel by transillumination at 302 nm.

### Immunizations

The native Enc and HEnc particles were immunized in 7 to 14 week-old, pathogen-free BALB/c and C57BL/6 inbred mice. Mice were ordered from Charles River Laboratories (Wilmington, MA) and housed in the Physiological Research Laboratory at Georgia Institute of Technology. All animal protocols and procedures were approved by the Georgia Tech Institutional Animal Care and Use Committee. PNPs were filter sterilized and resuspended to 0.5 mg/mL final concentrations in PBS. Immunizations were performed by injecting a total of 100 µL of PNP stock subcutaneously into the lower anterior abdomen (two 50 µL injections, one in each side of the abdomen). Body mass was noted and plotted over time for all groups. Serum samples were collected by submandibular bleeds every week (except week 1) to track serum antibody responses over time.

### Immune response assessment (ELISA)

Anti-scaffold total IgG and K417N peptide response were analysed using ELISA assays. Purified encapsulins, RBD (K417N/ WT/ Omicron BA.1) (Sino Biologicals) or streptavidin (catalogue #434302; Invitrogen) were plated on half-area high-binding 96-well polystyrene plates (Corning) at 1 µg/mL (in PBS) and stored at 4 °C overnight. The next day, the plates were washed with PBST (1x PBS with 0.5% Tween), then blocked with 1% casein (w/v) in PBS (VWR International, Radnor, PA) for 2 hours at room temperature with mild shaking (55 rpm). Except for the peptide analysis, the streptavidin plates were washed after 1 hour and coated with 5 µg/ml of Biotin-PEG_4_-K417N peptide, then incubated for another hour. Serum antibody titers were calculated by serially diluting the serum samples in block buffer (from 1:125 to 1:512,000) and plating the resulting solutions in duplicate after washing away unbound peptides. Following a 1-hour incubation, secondary reporter goat anti-mouse IgG HRP (Southern Biotech) was diluted (1:2500 for encapsulins, RBDs and 1:2000 for peptides) in blocking buffer, added to the plates, and incubated for 1 hour at room temperature with shaking. The plates were washed one final time with PBST and were then developed by adding 1-step Ultra TMB (Fisher Scientific) for 60 seconds, followed by quenching with 2N H_2_SO_4_. Absorbance values at 450 nm were measured using a plate reader (Varioskan Flash, Thermo Fisher Scientific). Antibody titers were calculated by plotting the serum dilution values (log scale) versus the absorbance values at 450 nm, fitting the datasets to sigmoidal non-linear regression curves, and extracting the resulting IC_50_ values from the sigmoidal plots (all analyses performed using GraphPad Prism v.8).

### Serum Biolayer Interferometry (BLI) assay

Analysis of serum antibodies binding to PNP scaffolds by biolayer interferometry (BLI) was performed on an Octet R4 (Sartorius) at 30 °C in kinetics buffer (PBS, 0.05% BSA, 0.02% Tween20) using Greiner F-bottom medium binding 96 well plates (catalogue #655076). Biosensors were equilibrated for 10 minutes in kinetics buffer and microplates filled with 200 µL of sample in kinetic buffer, then agitated at 1000 rpm. Anti-mouse Fc (AMC) biosensors were loaded for 300 seconds with serum IgG (serum dilution 1:200) giving a loading response of 0.9 ± 0.07 nm. A baseline was collected for 120 seconds to demonstrate stable loading. The loaded sensors were then allowed to associate with either native Enc (WT) or PP7-PP7 (WT) protein particles at 30 µg/mL for 300 seconds and dissociated in kinetic buffer for 1800 seconds. Collected data were analyzed using the Octet Data Analysis studio software. The buffer-subtracted Octet data were fit locally (kinetic analysis) to a simple 1:1 Langmuir model, all traces were aligned to the start of the association step, and inter-step corrections were applied to align all the steps. Since no appreciable disassociation was observed between the serum antibodies and the PNPs, K_D_ (apparent equilibrium dissociation constant) could not be calculated, thus only the *k*_a_ (association rate constant) values are reported. Kinetic curves are also reported for reference (average curve fit R^2^ values > 0.95).

### Immune response assessment (ELISpot)

For B cell frequency assessment, the highest responding mice were selected based on their serum titers (two from each group) and dosed with 25 µg/mice of the Enc-K417N variants 5 days before analysis. All the following steps were done under sterile condition. The day before harvest, ELISpot plates (catalogue #MSIPS4510, Millipore) were activated for 60 seconds with 15 µl/well of 35% ethanol in water, washed 3x with PBS, and then coated with filter-sterilized antigens (PP7-PP7, PP7-K417N-PP7, Enc variants or RBD (K417N/ WT/ Omicron BA.1) in PBS. Plates were incubated at 4 °C overnight, then washed with PBS and then blocked with RPMI (RPMI-1640, 10% fetal bovine serum, 1% non-essential amino acids, penicillin, streptomycin, glutamine) media for 2 hrs at 37 °C incubator. Mouse spleens were harvested, and single cell suspensions were generated by passing them through a 70 µm nylon filter resuspended in RPMI media. Cells were spun at 300*g* for 8 mins and the red blood cells in the suspension were lysed by adding RBC lysis buffer (2 mL/spleen; catalogue #R7767, Sigma) and incubated on ice for 3-5 mins until the solution appeared clear. The lysis was quenched by adding RPMI and removed by centrifugation and pelleting the cells. Cells were resuspended in RPMI and counted to make up to a 5 x 10^6^ cells/mL concentration. 1 million cells were plated in triplicate for PP7 (WT), PP7-K417N-PP7, and RBD antigen-containing wells, while 0.5 million cells were plated in triplicated for Enc variant antigen wells. Plates were incubated in a cell culture incubator at 37 °C, 5% CO_2_ and 97% humidity in the dark for 48 hours without disturbance. The plates were then washed 5x times with PBST to remove cells and debris, then incubated at room temperature with biotinylated secondary goat anti-mouse IgG (catalogue #3825-6-250, Mabtech) at a 2 µg/mL concentration in the dark for 2 hours. Plates were washed 6x times with PBST and plated with streptavidin-HRP conjugate (1:1000 dilution; catalogue #3310-9-1000, Mabtech) for 1 hour at room temperature in the dark. The plates were then washed 3x with PBST, 3x with PBS, then followed by addition of 100 µL/well of ELISpot substrate (catalogue #3651-10; Mabtech). Plates were incubated in the dark for up to 25 minutes while the spot formation was closely monitored. The reaction was quenched by running the plate under running tap water.

For T cell ELISpot, the protocol was similar to the B cell ELISpot with the following differences: the day before harvest, plates were coated with anti-mouse IFN-γ (catalogue #14731385, Invitrogen) and anti-mouse IL-4 (catalogue #14704185, Invitrogen). Stimulating antigens (filter sterilized Enc particles at 1 µg/mL), Concanavalin A (as positive control at 1 µg/mL), or PBS (as negative control) were added along with the cells at 0.5 million cells per well. Secondary antibodies biotinylated rat anti-mouse-IFN-γ (catalogue #13704285, Invitrogen) and biotinylated rat anti-mouse-IL4 (catalogue #13731281, Invitrogen) were used. Plates were stored in the dark under dry conditions until ready to be read.

### Flow cytometry analysis

BALB/c mice were immunized with three-doses at three weeks apart with either PBS, homologous or heterologous approaches by subcutaneous injections in the anterior lower abdomen around the inguinal mammary fat pads (50uL on both sides; 50ug of particle). Two weeks after final boost, serum was collected by cardiac puncture for terminal serum response, and lymphoid organs were harvested for flow cytometry. Both right and left inguinal lymph nodes and the spleen were extracted from each mouse and placed in PBS on ice. The spleen and lymph nodes were mashed into 70um mesh size EASYstrainer Cell Strainers (Greiner Bio-One 542070) and washed with PBS. The recovered cells in PBS were spun down at 400xg for 5 minutes and the supernatant was disposed. For spleen samples, red blood cell lysis was performed by adding 1mL of Red Blood Cell Lysing Buffer Hybri-Max™ (Sigma R7757) per spleen sample cell pellet, letting that incubate at room temperature for 12 minutes, and then quenching the samples with 30mL of PBS. These samples were again spun down at 400g for 5 mins and the supernatant was disposed. The lymph node cell pellets were resuspended in 200uL PBS, and the red blood cell lysed spleen cell pellets were resuspended in 10mL PBS. Then 200uL of each of these samples were plated in 96 well U-bottom Tissue Culture Plates (Falcon 353227). The plate was spun down and the supernatant was disposed. Anti-mouse CD16/CD32 Fc Block antibody purchased at a concentration of 2mg/mL (Tonbo Biosciences 40-0161-M001) was added at 0.5 uL per sample (in 100uL total volume per sample of block antibody diluted in PBS) to the samples in the well plate, and incubated on ice for 5 minutes. The plate was spun down and the supernatant was disposed. All samples were washed once with PBS and then prepared for staining. Lyophilized Zombie Aqua™ dye (Biolegend 423102) was resuspended in 100uL DMSO, and 0.5uL of this solution was added per sample, and incubated in the dark at room temperature for 30 minutes. The plate was spun down and washed once with FACS buffer (0.5% BSA in PBS; Sigma), then resuspended in 100ul of FACS buffer.

To stain K417N-peptide reactive B-cells, 0.25 mg of streptavidin was labelled with 0.45 umol Alexa-Fluor 647 at room temperature for 2 hours and cleaning up the reaction with PD-10 column and four rounds of buffer exchange with spin filtration. The biotin-K417N peptide (VRQIAPGQTG**<u>N</u>**IADYNYKLP ; GenScript) was mixed with the streptavidin-AF647 in a 4:1 ratio at room temperature for 2 hours. Free D-biotin (10ug) was then added to occupy any remaining streptavidin binding sites not already occupied with biotin-K417N peptide and purified by Amicon® Ultra Centrifugal Filter, 30 kDa MWCO (Sigma UFC9030). The AF647-streptavidin-biotin-K417N peptide solution was diluted to a concentration of 0.036mg/mL in PBS, and incubated with 100uL per sample (total of 3.6ug antigen per sample) in the dark for 25 minutes. The plate was spun down and the supernatant was disposed. All samples were washed once with 100 ul FACS buffer and resuspended in 100 ul of master mix per sample. Stock master mix was prepared as followed: All antibodies were purchased at stock concentration of 0.2mg/mL. Following volumes were added in a single tube containing 100ul FACS buffer per sample: 0.625uL per sample of APC/Cyanine7 anti-mouse CD19 (Biolegend 152412), 1.25uL per sample of Brilliant Violet 421™ anti-mouse CD38 (Biolegend 102732), and 1.25uL per sample of Brilliant Violet 605™ anti-mouse CD95/FAS (Biolegend 152612). Samples were incubated with the master mix in the dark on ice for 15 minutes. The plate was spun down, the supernatant disposed and then washed once with 100 ul of FACS buffer. The cells were resuspended in 100uL FACS buffer and placed in the Cytek Aurora 5 Laser flow cytometer. The flow cytometer recorded 85uL per sample, recording cells stained with AF647, BV421, BV605, APC/Cy7, and Zombie Aqua dyes.

Single stain reference controls for all markers except peptide were performed on splenocytes collected from naïve mouse. The spleen was prepared as described above and resuspended in 10mL. Then 200uL of this splenocyte solution per well was plated for the CD19, CD38, and CD95 antibody controls. Additionally, 200uL of heat-killed splenocyte sample (at 45°C for 10 minutes) was plated as negative population for Zombie Aqua staining. Cells were stained as described above, except with only one antibody stain per sample. For the AF647-streptavidin-biotin-K417N peptide single stain control, splenocytes from the approach (2; PP7-HEnc-PP7) with the highest ELISA anti-K417N titer were used as the single stain control sample. Data analysis was performed with FlowJo software (FlowJo, LLC) and data plotted using GraphPad Prism v10.

## Acknowledgements

This work was supported by the Centers for Disease Control and Prevention (75D30121P10377), the NIH (R21 AI166639), and by the Simons Foundation (SF349247) for structural studies at the Simons Electron Microscopy Center at the New York Structural Biology Center. Figures 2A, 3A,B, 5A, 6A, 7A and the TOC graphic were created partly using BioRender.com (http://biorender.com/).

