## Supplemental Material for "Heterologous Prime-Boost with Immunologically Orthogonal Protein Nanoparticles for Peptide Immunofocusing"

### Supporting Information

### Plasmid MCS sequences of encapsulin variants

**Bold text** = T7 promoter/terminator sequences
Yellow highlight = ribosomal binding site
Light blue highlight = Shot47sss aptamer sequence
Dark yellow highlight = K417N peptide sequence
**Bold and underlined text** = Translation start codon
Red highlight = His$_6$ affinity tag
Green highlight = encapsulin coat protein

pET23b:Enc

**TAATACGACTCACTATAGG**GAGACCACAACGGTTTCCCTCTAGAAATAATTTTGTTTAACTTTAAGAAGGAGATATA
CAT**ATG**CCGGATTTCCTCGGTCATGCGGAAAATCCGCTGCGTGAGGAAGAATGGGCCCGTCTTAACGAAACGGTCAT
TCAGGTCGCTCGTCGCAGTCTGGTTGGTCGTCGGATTCTGGACATTTACGGGCCATTAGGTGCCGGAGTACAAACCG
TCCCGTATGACGAGTTTCAGGGCGTTAGCCCTGGTGCTGTCGATATTGTGGGTGAACAGGAAACTGCGATGGTGTTT
ACGGATGCCCGTAAATTCAAGACCATTCCCATCATCTACAAAGACTTCCTGCTGCATTGGCGGGATATCGAAGCAGC
ACGCACACACAACATGCCGTTAGACGTATCGGCAGCAGCAGGGGCCGCTGCGCTTTGTGCTCAACAGGAGGACGAAC
TGATTTTCTATGGCGATGCGCGTTTGGGGTATGAAGGCCTGATGACCGCCAATGGTCGCCTGACCGTGCCTCTGGGA
GATTGGACCAGTCCGGGTGGCGGCTTTCAGGCGATTGTGGAAGCCACGCGCAAACTGAATGAGCAAGGCCACTTTGG
TCCGTATGCGGTTGTGTTATCGCCACGCCTGTACTCCCAATTGCATCGCATCTACGAGAAAACGGGCGTTCTGGAGA
TTGAAACCATTCGCCAGTTAGCATCAGATGGCGTGTATCAGAGCAATCGTCTGCGTGGCGAATCTGGCGTCGTGGTA
TCTACAGGGCGCGAGAACATGGATCTTGCCGTTTCTATGGACATGGTTGCGGCTTATCTGGGCGCGAGCCGCATGAA
CCATCCGTTTCGCGTGTTGGAAGCGCTCTTGCTCCGCATCAAACACCCCGATGCGATCTGCACTCTGGAAGGAGCCG
GTGCGACCGAACGCCGTTAAAAGCTTGCGGCCGCACTCGAGCACCACCACCACCACCACTGAGATCCGGCTGCTAAC
AAAGCCCGAAAGGAAGCTGAGTTGGCTGCTGCCACCGCTGAGCAATAA**CTAGCATAACCCCTTGGGGCCTCTAAACG
GGTCTTGAGGGGTTTTTTG**

pET14b:HEnc

**TAATACGACTCACTATAGG**GAGACCACAACGGTTTCCCTCTAGAAATAATTTTGTTTAACTTTAAGAAGGAGATATA
CC**ATG**GGCAGCAGCCATCATCATCATCATCACAGCAGCGGCCTGGTGCCGCGCGGCAGCCATATGCCGGATTTCCTC
GGTCATGCGGAAAATCCGCTGCGTGAGGAAGAATGGGCCCGTCTTAACGAAACGGTCATTCAGGTCGCTCGTCGCAG
TCTGGTTGGTCGTCGGATTCTGGACATTTACGGGCCATTAGGTGCCGGAGTACAAACCGTCCCGTATGACGAGTTTC
AGGGCGTTAGCCCTGGTGCTGTCGATATTGTGGGTGAACAGGAAACTGCGATGGTGTTTACGGATGCCCGTAAATTC
AAGACCATTCCCATCATCTACAAAGACTTCCTGCTGCATTGGCGGGATATCGAAGCAGCACGCACACACAACATGCC
GTTAGACGTATCGGCAGCAGCAGGGGCCGCTGCGCTTTGTGCTCAACAGGAGGACGAACTGATTTTCTATGGCGATG
CGCGTTTGGGGTATGAAGGCCTGATGACCGCCAATGGTCGCCTGACCGTGCCTCTGGGAGATTGGACCAGTCCGGGT
GGCGGCTTTCAGGCGATTGTGGAAGCCACGCGCAAACTGAATGAGCAAGGCCACTTTGGTCCGTATGCGGTTGTGTT
ATCGCCACGCCTGTACTCCCAATTGCATCGCATCTACGAGAAAACGGGCGTTCTGGAGATTGAAACCATTCGCCAGT
TAGCATCAGATGGCGTGTATCAGAGCAATCGTCTGCGTGGCGAATCTGGCGTCGTGGTATCTACAGGGCGCGAGAAC
ATGGATCTTGCCGTTTCTATGGACATGGTTGCGGCTTATCTGGGCGCGAGCCGCATGAACCATCCGTTTCGCGTGTT
GGAAGCGCTCTTGCTCCGCATCAAACACCCCGATGCGATCTGCACTCTGGAAGGAGCCGGTGCGACCGAACGCCGTT
AAGGATCCTATTGCTAACAAAGCCCGAAAGGAAGCTGAGTTGGCTGCTGCCACCGCTGAGCAATAA**CTAGCATAACC
CCTTGGGGCCTCTAAACGGGTCTTGAGGGGTTTTTTG**

pET14b:*HEnc

**TAATACGACTCACTATAGG**GAGACCACAACGGTTTCCCGGGTATATTGGCGCCTTCGTGGAATGTCAGTGCCTGGTC
TAGAAATAATTTTGTTTAACTTTAAGAAGGAGATATACC**ATG**GGCAGCAGCCATCATCATCATCATCACAGCAGCGG
CCTGGTGCCGCGCGGCAGCCATATGCCGGATTTCCTCGGTCATGCGGAAAATCCGCTGCGTGAGGAAGAATGGGCCC
GTCTTAACGAAACGGTCATTCAGGTCGCTCGTCGCAGTCTGGTTGGTCGTCGGATTCTGGACATTTACGGGCCATTA
GGTGCCGGAGTACAAACCGTCCCGTATGACGAGTTTCAGGGCGTTAGCCCTGGTGCTGTCGATATTGTGGGTGAACA
GGAAACTGCGATGGTGTTTACGGATGCCCGTAAATTCAAGACCATTCCCATCATCTACAAAGACTTCCTGCTGCATT
GGCGGGATATCGAAGCAGCACGCACACACAACATGCCGTTAGACGTATCGGCAGCAGCAGGGGCCGCTGCGCTTTGT

GCTCAACAGGAGGACGAACTGATTTTCTATGGCGATGCGCGTTTGGGGTATGAAGGCCTGATGACCGCCAATGGTCG
CCTGACCGTGCCTCTGGGAGATTGGACCAGTCCGGGTGGCGGCTTTCAGGCGATTGTGGAAGCCACGCGCAAACTGA
ATGAGCAAGGCCACTTTGGTCCGTATGCGGTTGTGTTATCGCCACGCCTGTACTCCCAATTGCATCGCATCTACGAG
AAAACGGGCGTTCTGGAGATTGAAACCATTCGCCAGTTAGCATCAGATGGCGTGTATCAGAGCAATCGTCTGCGTGG
CGAATCTGGCGTCGTGGTATCTACAGGGCGCGAGAACATGGATCTTGCCGTTTCTATGGACATGGTTGCGGCTTATC
TGGGCGCGAGCCGCATGAACCATCCGTTTCGCGTGTTGGAAGCGCTCTTGCTCCGCATCAAACACCCCGATGCGATC
TGCACTCTGGAAGGAGCCGGTGCGACCGAACGCCGTTAAGGATCCTATTGCTAACAAAGCCCGAAAGGAAGCTGAGT
TGGCTGCTGCCACCGCTGAGCAATAA**CTAGCATAACCCCTTGGGGCCTCTAAACGGGTCTTGAGGGGTTTTTTG**

pET14b:*HEnc*

**TAATACGACTCACTATAGG**GAGACCACAACGGTTTCCCGGGTATATTGGCGCCTTCGTGGAATGTCAGTGCCTGGTC
TAGAAATAATTTTGTTTAACTTTAAGAAGGAGATATACC**ATG**GGCAGCAGCCATCATCATCATCATCACAGCAGCGG
CCTGGTGCCGCGCGGCAGCCATATGCCGGATTTCCTCGGTCATGCGGAAAATCCGCTGCGTGAGGAAGAATGGGCCC
GTCTTAACGAAACGGTCATTCAGGTCGCTCGTCGCAGTCTGGTTGGTCGTCGGATTCTGGACATTTACGGGCCATTA
GGTGCCGGAGTACAAACCGTCCCGTATGACGAGTTTCAGGGCGTTAGCCCTGGTGCTGTCGATATTGTGGGTGAACA
GGAAACTGCGATGGTGTTTACGGATGCCCGTAAATTCAAGACCATTCCCATCATCTACAAAGACTTCCTGCTGCATT
GGCGGGATATCGAAGCAGCACGCACACACAACATGCCGTTAGACGTATCGGCAGCAGCAGGGGCCGCTGCGCTTTGT
GCTCAACAGGAGGACGAACTGATTTTCTATGGCGATGCGCGTTTGGGGTATGAAGGCCTGATGACCGCCAATGGTCG
CCTGACCGTGCCTCTGGGAGATTGGACCAGTCCGGGTGGCGGCTTTCAGGCGATTGTGGAAGCCACGCGCAAACTGA
ATGAGCAAGGCCACTTTGGTCCGTATGCGGTTGTGTTATCGCCACGCCTGTACTCCCAATTGCATCGCATCTACGAG
AAAACGGGCGTTCTGGAGATTGAAACCATTCGCCAGTTAGCATCAGATGGCGTGTATCAGAGCAATCGTCTGCGTGG
CGAATCTGGCGTCGTGGTATCTACAGGGCGCGAGAACATGGATCTTGCCGTTTCTATGGACATGGTTGCGGCTTATC
TGGGCGCGAGCCGCATGAACCATCCGTTTCGCGTGTTGGAAGCGCTCTTGCTCCGCATCAAACACCCCGATGCGATC
TGCACTCTGGAAGGAGCCGGTGCGACCGAACGCCGTTAAGGATCCTATTGCTAAGGGTATATTGGCGCCTTCGTGGA
ATGTCAGTGCCTGGGCTGAGCAATAA**CTAGCATAACCCCTTGGGGCCTCTAAACGGGTCTTGAGGGGTTTTTTG**

pET14b:HEnc-K417$N_A$

**TAATACGACTCACTATAGG**GAGACCACAACGGTTTCCCTCTAGAAATAATTTTGTTTAACTTTAAGAAGGAGATATA
CC**ATG**GGCAGCAGCCATCATCATCATCATCACAGCAGCGGCCTGGTGCCGCGCGGCAGCCATATGCCGGATTTCCTC
GGTCATGCGGAAAATCCGCTGCGTGAGGAAGAATGGGCCCGTCTTAACGAAACGGTCATTCAGGTCGCTCGTCGCAG
TCTGGTTGGTCGTCGGATTCTGGACATTTACGGGCCATTAGGTGCCGGAGTACAAACCGTCCCGTATGACGAGTTTC
AGGGCGTGCGCCAGATTGCACCAGGTCAGACCGGTAACATCGCGGATTATAACTACAAACTGCCGGTTAGCCCTGGT
GCTGTCGATATTGTGGGTGAACAGGAAACTGCGATGGTGTTTACGGATGCCCGTAAATTCAAGACCATTCCCATCAT
CTACAAAGACTTCCTGCTGCATTGGCGGGATATCGAAGCAGCACGCACACACAACATGCCGTTAGACGTATCGGCAG
CAGCAGGGGCCGCTGCGCTTTGTGCTCAACAGGAGGACGAACTGATTTTCTATGGCGATGCGCGTTTGGGGTATGAA
GGCCTGATGACCGCCAATGGTCGCCTGACCGTGCCTCTGGGAGATTGGACCAGTCCGGGTGGCGGCTTTCAGGCGAT
TGTGGAAGCCACGCGCAAACTGAATGAGCAAGGCCACTTTGGTCCGTATGCGGTTGTGTTATCGCCACGCCTGTACT
CCCAATTGCATCGCATCTACGAGAAAACGGGCGTTCTGGAGATTGAAACCATTCGCCAGTTAGCATCAGATGGCGTG
TATCAGAGCAATCGTCTGCGTGGCGAATCTGGCGTCGTGGTATCTACAGGGCGCGAGAACATGGATCTTGCCGTTTC
TATGGACATGGTTGCGGCTTATCTGGGCGCGAGCCGCATGAACCATCCGTTTCGCGTGTTGGAAGCGCTCTTGCTCC
GCATCAAACACCCCGATGCGATCTGCACTCTGGAAGGAGCCGGTGCGACCGAACGCCGTTAAGGATCCTATTGCTAA
CAAAGCCCGAAAGGAAGCTGAGTTGGCTGCTGCCACCGCTGAGCAATAA**CTAGCATAACCCCTTGGGGCCTCTAAAC**
**GGGTCTTGAGGGGTTTTTTG**

pET14b:HEnc-K417N$_B$

**TAATACGACTCACTATAGG**GAGACCACAACGGTTTCCCTCTAGAAATAATTTTGTTTAACTTTAAGAAGGAGATATA
CC**<u>ATG</u>**GGCAGCAGCCATCATCATCATCATCACAGCAGCGGCCTGGTGCCGCGCGGCAGCCATATGCCGGATTTCCTC
GGTCATGCGGAAAATCCGCTGCGTGAGGAAGAATGGGCCCGTCTTAACGAAACGGTCATTCAGGTCGCTCGTCGCAG
TCTGGTTGGTCGTCGGATTCTGGACATTTACGGGCCATTAGGTGCCGGAGTACAAACCGTCCCGTATGACGAGTTTC
AGGGCGTTAGCCCTGGTGCTGTCGATATTGTGGGTGAACAGGAAACTGCGATGGTGTTTACGGATGCCCGTAAATTC
AAGACCATTCCCATCATCTACAAAGACTTCCTGCTGCATTGGCGGGATATCGAAGCAGCACGCACACACAACATGCC
GTTAGACGTATCGGCAGCAGCAGGGGCCGCTGCGCTTTGTGCTCAACAGGAGGACGAACTGATTTTCTATGGCGATG
CGCGTGTGCGCCAGATTGCACCAGGTCAGACCGGTAACATCGCGGATTATAACTACAAACTGCCGTTGGGGTATGAA
GGCCTGATGACCGCCAATGGTCGCCTGACCGTGCCTCTGGGAGATTGGACCAGTCCGGGTGGCGGCTTTCAGGCGAT
TGTGGAAGCCACGCGCAAACTGAATGAGCAAGGCCACTTTGGTCCGTATGCGGTTGTGTTATCGCCACGCCTGTACT
CCCAATTGCATCGCATCTACGAGAAAACGGGCGTTCTGGAGATTGAAACCATTCGCCAGTTAGCATCAGATGGCGTG
TATCAGAGCAATCGTCTGCGTGGCGAATCTGGCGTCGTGGTATCTACAGGGCGCGAGAACATGGATCTTGCCGTTTC
TATGGACATGGTTGCGGCTTATCTGGGCGCGAGCCGCATGAACCATCCGTTTCGCGTGTTGGAAGCGCTCTTGCTCC
GCATCAAACACCCCGATGCGATCTGCACTCTGGAAGGAGCCGGTGCGACCGAACGCCGTTAAGGATCCTATTGCTAA
CAAAGCCCGAAAGGAAGCTGAGTTGGCTGCTGCCACCGCTGAGCAATAA**CTAGCATAACCCCTTGGGGCCTCTAAAC**
**GGGTCTTGAGGGGTTTTTTG**

pET14b:*HEnc*-K417N$_B$

**TAATACGACTCACTATAGG**GAGACCACAACGGTTTCCCGGGTATATTGGCGCCTTCGTGGAATGTCAGTGCCTGGTC
TAGAAATAATTTTGTTTAACTTTAAGAAGGAGATATACC**<u>ATG</u>**GGCAGCAGCCATCATCATCATCATCACAGCAGCGG
CCTGGTGCCGCGCGGCAGCCATATGCCGGATTTCCTCGGTCATGCGGAAAATCCGCTGCGTGAGGAAGAATGGGCCC
GTCTTAACGAAACGGTCATTCAGGTCGCTCGTCGCAGTCTGGTTGGTCGTCGGATTCTGGACATTTACGGGCCATTA
GGTGCCGGAGTACAAACCGTCCCGTATGACGAGTTTCAGGGCGTTAGCCCTGGTGCTGTCGATATTGTGGGTGAACA
GGAAACTGCGATGGTGTTTACGGATGCCCGTAAATTCAAGACCATTCCCATCATCTACAAAGACTTCCTGCTGCATT
GGCGGGATATCGAAGCAGCACGCACACACAACATGCCGTTAGACGTATCGGCAGCAGCAGGGGCCGCTGCGCTTTGT
GCTCAACAGGAGGACGAACTGATTTTCTATGGCGATGCGCGTGTGCGCCAGATTGCACCAGGTCAGACCGGTAACAT
CGCGGATTATAACTACAAACTGCCGTTGGGGTATGAAGGCCTGATGACCGCCAATGGTCGCCTGACCGTGCCTCTGG
GAGATTGGACCAGTCCGGGTGGCGGCTTTCAGGCGATTGTGGAAGCCACGCGCAAACTGAATGAGCAAGGCCACTTT
GGTCCGTATGCGGTTGTGTTATCGCCACGCCTGTACTCCCAATTGCATCGCATCTACGAGAAAACGGGCGTTCTGGA
GATTGAAACCATTCGCCAGTTAGCATCAGATGGCGTGTATCAGAGCAATCGTCTGCGTGGCGAATCTGGCGTCGTGG
TATCTACAGGGCGCGAGAACATGGATCTTGCCGTTTCTATGGACATGGTTGCGGCTTATCTGGGCGCGAGCCGCATG
AACCATCCGTTTCGCGTGTTGGAAGCGCTCTTGCTCCGCATCAAACACCCCGATGCGATCTGCACTCTGGAAGGAGC
CGGTGCGACCGAACGCCGTTAAGGATCCTATTGCTAAGGGTATATTGGCGCCTTCGTGGAATGTCAGTGCCTGGGCT
GAGCAATAA**CTAGCATAACCCCTTGGGGCCTCTAAACGGGTCTTGAGGGGTTTTTTG**

### Plasmid MCS sequences of PP7-PP7 variants

**Bold text** = T7 promoter/terminator sequences
Yellow highlight = ribosomal binding site
Green highlight = PP7 N-terminal coat protein
Magenta highlight = PP7 N-terminal coat protein
**Bold and underlined text** = Translation start codon
Dark yellow highlight = K417N peptide sequence
Grey highlight = AYGG linker

pCDF:PP7-PP7

**TAATACGACTCACTATAGGGG**GGAATTGTGAGCGGATAACAATTCCCCTGTAGAAATAATTTTGTTTAACTTTAAGA
AGGAGATATAGGATCC**ATG**AGCAAAACCATTGTTCTGAGCGTGGGTGAAGCGACCCGTACCCTGACCGAAATCCAGA
GCACCGCTGACCGTCAAATTTTTGAGGAAAAAGTGGGTCCGCTGGTTGGCCGTCTGCGTCTGACCGCGAGCCTGCGT
CAGAACGGTGCGAAGACCGCGTACCGTGTGAACCTGAAACTGGACCAAGCGGATGTGGTTGATTGCAGCACCAGCGT
TTGCGGCGAGCTGCCGAAAGTGCGTTACACCCAGGTTTGGAGCCACGATGTGACCATCGTTGCGAACAGCACCGAAG
CGAGCCGTAAGAGCCTGTATGATCTGACCAAAAGCCTGGTGGCGACCAGCCAAGTTGAGGACCTGGTGGTTAATCTT
GTACCACTTGGTCGCGCATATGGCGGTTCGAAAACCATCGTCCTGTCCGTGGGCGAAGCAACCCGCACCCTGACCGA
AATCCAATCTACCGCAGACCGCCAAATCTTTGAAGAAAAAGTGGGTCCGCTGGTCGGTCGTCTGCGTCTGACCGCCT
CTCTGCGTCAGAACGGCGCGAAAACGGCCTATCGCGTCAATCTGAAACTGGATCAAGCAGACGTGGTTGATTGCAGC
ACCTCTGTTTGTGGTGAACTGCCGAAAGTGCGTTATACGCAGGTTTGGTCACATGACGTCACCATTGTGGCAAACTC
GACGGAAGCTAGTCGCAAATCCCTGTACGATCTGACCAAATCCCTGGTGGCGACCTCTCAAGTGGAAGACCTGGTGG
TGAACCTGGTGCCGCTGGGCCGCTAACTCGAGTCTGGTAAAGAAACCGCTGCTGCGAAATTTGAACGCCAGCACATG
GACTCGTCTACTAGCGCAGCTTAATTAACCTAGGCTGCTGCCACCGCTGAGCAATAAC**TAGCATAACCCCTTGGGGC**
**CTCTAAACGGGTCTTGAGGGGTTTTTTG**

pCDF:PP7-K417N-PP7

**TAATACGACTCACTATAGGGG**GGAATTGTGAGCGGATAACAATTCCCCTGTAGAAATAATTTTGTTTAACTTTAAGA
AGGAGATATAGGATCC**ATG**AGCAAAACCATTGTTCTGAGCGTGGGTGAAGCGACCCGTACCCTGACCGAAATCCAGA
GCACCGCTGACCGTCAAATTTTTGAGGAAAAAGTGGGTCCGCTGGTTGGCCGTCTGCGTCTGACCGCGAGCCTGCGT
CAGAACGGTGCGAAGACCGCGTACCGTGTGAACCTGAAACTGGACCAAGCGGATGTGGTTGATTGCAGCACCAGCGT
TTGCGGCGAGCTGCCGAAAGTGCGTTACACCCAGGTTTGGAGCCACGATGTGACCATCGTTGCGAACAGCACCGAAG
CGAGCCGTAAGAGCCTGTATGATCTGACCAAAAGCCTGGTGGCGACCAGCCAAGTTGAGGACCTGGTGGTTAATCTT
GTACCACTTGGTCGCGCATATGGCGTGCGTCAGATCGCTCCGGGCCAGACCGGGAACATTGCGGACTACAACTACAA
GTTACCCGGTTCGAAAACCATCGTCCTGTCCGTGGGCGAAGCAACCCGCACCCTGACCGAAATCCAATCTACCGCAG
ACCGCCAAATCTTTGAAGAAAAAGTGGGTCCGCTGGTCGGTCGTCTGCGTCTGACCGCCTCTCTGCGTCAGAACGGC
GCGAAAACGGCCTATCGCGTCAATCTGAAACTGGATCAAGCAGACGTGGTTGATTGCAGCACCTCTGTTTGTGGTGA
ACTGCCGAAAGTGCGTTATACGCAGGTTTGGTCACATGACGTCACCATTGTGGCAAACTCGACGGAAGCTAGTCGCA
AATCCCTGTACGATCTGACCAAATCCCTGGTGGCGACCTCTCAAGTGGAAGACCTGGTGGTGAACCTGGTGCCGCTG
GGCCGCTAACTCGAGTCTGGTAAAGAAACCGCTGCTGCGAAATTTGAACGCCAGCACATGGACTCGTCTACTAGCGC
AGCTTAATTAACCTAGGCTGCTGCCACCGCTGAGCAATAAC**TAGCATAACCCCTTGGGGCCTCTAAACGGGTCTTGA**
**GGGGTTTTTTG**

**Preparation of loop-inserted *M. xanthus* Enc particles**

*Insertion point C in Enc coat protein.* It is noted in the main text that no particles were isolated from the protein bearing the 20-amino-acid insertion at between residues N145 and G146 ($IP_C$). This was a surprising result as the structurally homologous position in the extensively studied encapsulin from *Thermotoga maritima* (i.e., between residues 138/139) has been highly tolerant of peptide insertions.[1-3] One possible explanation for this difference is the expected greater stability of the promoter fold in the hyperthermophilic *T. maritima* encapsulin relative to the mesophilic *M. xanthus* container. Indeed, recent efforts to study the disassembly and reassembly behavior of several encapsulin variants showed that the *M. xanthus* encapsulins were both less thermostable and less tolerant of chemical denaturants than those from *T. maritima*.[4]

**RT-qPCR analysis**

In order to accurately assess the levels of encapsulin protomer mRNAs specifically entrained within PNPs relative to non-specifically packaged host *E. coli* RNAs, two separate primer sets were generated for TaqMan-based RT-qPCR analyses (Table 1). The first primer set (primers 606/MG1655/F, 607/MG1655/R, and 608/MG1655/probe) were designed to target mRNAs generated from the idnT gene encoded in the host *E. coli* genome. The second set of primers (primers 609/MxEncA/F, 610/MxEncA/R, and 611/MxEncA/probe) were designed to target encapsulin protomer mRNAs in the various encapsulin plasmids. Standard curves utilizing these two primer sets were employed to quantify mRNA levels from purified PNP samples (*vide infra*).

Prior to RT-qPCR analyses, purified encapsulin samples were diluted to a final concentration of 1 mg/mL in 200 µL of 1x DNase I Buffer (New England Biolabs) and were treated with a final concentration of 20 U/mL of DNase I for 1 hour at 37 °C to eliminate any residual DNAs potentially bound to the encapsulins' exterior surfaces.[5] The DNase I was then heat inactivated at 75 °C for 10 minutes, and the treated PNP samples were transferred into reinforced 2 mL screw-top tubes with red silicone cap O-rings. Each screw-top tube was pre-aliquoted with 200 µL of TNA lysis buffer (Omega Bio-Tek, catalog #TNA-1000) and five 2.8 mm ceramic beads. The PNPs were then mechanically disrupted by bead mill homogenization for 45 seconds in an OMNI Bead Ruptor (OMNI International) operating at 6 m/s, followed by centrifugation for 3 minutes at 10,000 rpms. Extraction of liberated RNAs was then performed using the Omega Bio-Tek DNA/RNA kit (catalog #M6246-03) in conjunction with a Kingfisher Flex System (Thermo Fisher Scientific). Subsequently, a volume of 2 µL of each extracted sample was added in triplicate to separate wells of a 384-well plate using an ASSIST PLUS liquid handling robot (INTEGRA Biosciences). Next, a volume of 8 µL of master mix solution was added into each sample well using an Echo 525 liquid handler (Beckman Coulter). The master mix solution consisted of a 4 µL of RealPCR* RNA Master Mix (IDEXX Laboratories, catalog #99-56280), and 4 µL of a solution containing 900 nM of the appropriate amplification primers (i.e., either primers 606 and 607 or primers 609 and 610), and 250 nM of the probe primer (i.e., primer 608 or 611). RT-qPCR was then performed on either a QuantStudio™ 6 Flex or a QuantStudio™ 5 (Thermo Fisher Scientific) using the following protocol: reverse transcription at 50 °C for 15 minutes; initial denaturation at 95 °C for 1 minute; 45 cycles of denaturation at 95 °C for 15 seconds followed by extension at 60 °C for 30 seconds. Experimental results were analyzed using the Thermo Fisher cloud program by setting a threshold value at 10% of the intensity generated by the most concentrated sample from the respective standard curve plot. Both PNP-derived and standard curve sample

$C_t$ values were averaged from the triplicate reactions. Final RNA counts were derived from semi-log nonlinear regression analyses of standard curve plots generated by plotting log Concentration versus $C_t$ (Figure S6).

Standard curves for the idnT and encapsulin primer sets were generated from total cell RNAs and from plasmids, respectively, extracted from cultured *E. coli* cells. Briefly, single *E. coli* BL21(DE3) colonies containing either the Enc, HEnc, *HEnc, or *HEnc* plasmids were inoculated into 50 mL of 2YT media containing 0.1 mg/mL carbenicillin and were grown at 37 °C in a shaking incubator set to 250 rpms for 18-20 hours. The cell cultures were then divided into 10 mL aliquots, then the aliquots were centrifuged at 4500 rpms for 15 minutes in a Beckman Coulter Allegra X-30r tabletop centrifuge. The supernatants were decanted and the resulting cell pellets were stored at -80 °C until use. For the collection of total cellular RNAs, one 10 mL culture aliquot was thawed for each encapsulin variant, and RNAs were extracted from the aliquots using Omega Bio-Tek DNA/RNA kits. For the collection of plasmid DNAs, another 10 mL culture aliquot was thawed for each encapsulin variant, and plasmids were extracted using HiSpeed Plasmid Midi kits (Qiagen, catalog #12643). The concentrations of all extracted nucleic acid materials were measured using a QX200 Droplet Digital PCR machine (Bio-Rad), and all materials were stored at -80 °C until use. Standard curves were subsequently generated from extracted RNAs and plasmids by generating successive 3-fold dilutions of the respective templates for a total of 6 concentration points. The serially diluted template samples were included alongside the PNP-extracted RNAs in the 384-well plates used for RT-qPCR analyses.

| Primer | Sequence (5' → 3') |
|---|---|
| 606/MG1655/F | GAAATAAGGGTTTCCATCACCGTCC |
| 607/MG1655/R | GACCCTGGCTTCTGGCTATTTAAAG |
| 608/MG1655/probe | 6-FAM-ACGTTTCACCAACCGTCAGA-BHQ1 |
| 609/MxEncA/F | GTGTATCAGAGCAATCGTC |
| 610/MxEncA/R | CAACCATGTCCATAGAAAC |
| 611/MxEncA/probe | HEX-GTCGTGGTATCTACAGGGC-BHQ1 |

6-FAM = 6-carboxyfluorescein
HEX = hexachlorofluorescein
BHQ1 = Black Hole Quencher®-1

**Table 1:** List of primers employed for RT-qPCR analyses.

**Discussion of serum antibody titers over time for heterologous immunization strategies**

Examination of antibody titers over the course of the full vaccination schedules reveals several interesting trends. At week 2, BALB/c mice that received a primary dose of PP7-K417N-PP7 particles (or *HEnc* particles) displayed highly similar anti-K417N IgG titers, suggesting good biological replication of the data (Figure 6B, main text). Similarly, mice primed with HEnc particles also exhibited highly uniform, albeit lower responses. At week 3, the two vaccination groups that were primed with encapsulin but then boosted with PP7 (i.e., groups "3" and "4") showed the highest increases in IgG titers, though these increases were short lived. A second booster dose consisting of PP7 nanoparticles was sufficient to sustain the higher antibody levels (approach "3") whereas a second booster with HEnc particles led to a decrease in titer levels (approach "4"). Both groups subsequently showed a gradual decrease until the final booster dose at week 7, at which point the IgG titers increased higher than either homologous vaccination regimen, but lower overall than the other heterologous regimens in which the mice were primed with PP7 particles (i.e., groups “1” and “2”). In contrast, we saw a more general trend for the outbred CD-1 mice in which all of the heterologous vaccination strategies, with the exception of the approach “4”, yielded consistently higher anti-peptide K417N IgG titers that largely did not exhibit the gradual decreases over time that were observed in the BALB/c mice (Figure 7B, main text).

**Supplemental Figures:**

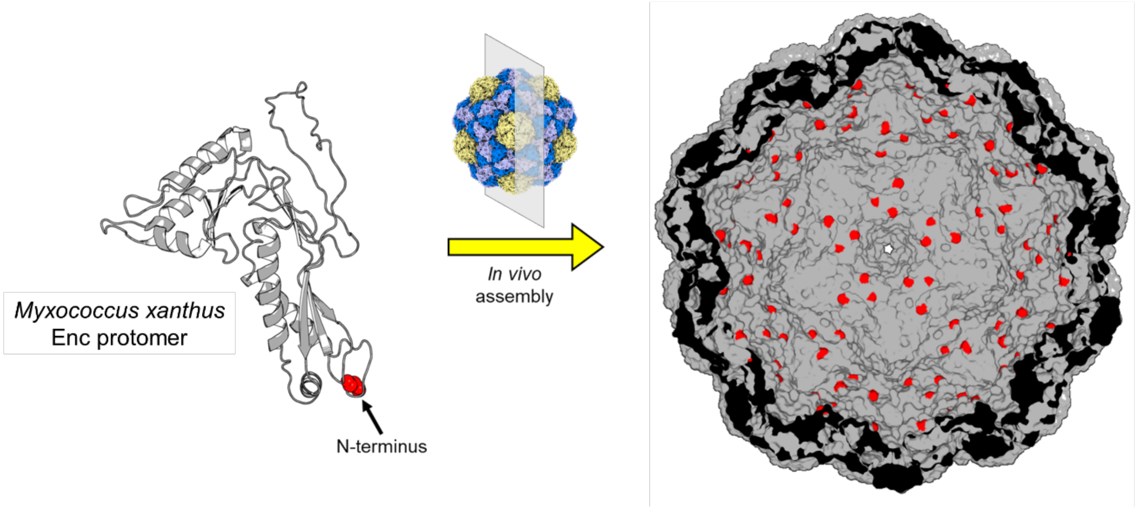

**Figure S1. Position of the N-terminus on encapsulin protomer** and the resulting luminal distribution of the N-termini in the assembled nanoparticles, as seen in a cross-sectional view (PDB: 7S2O).

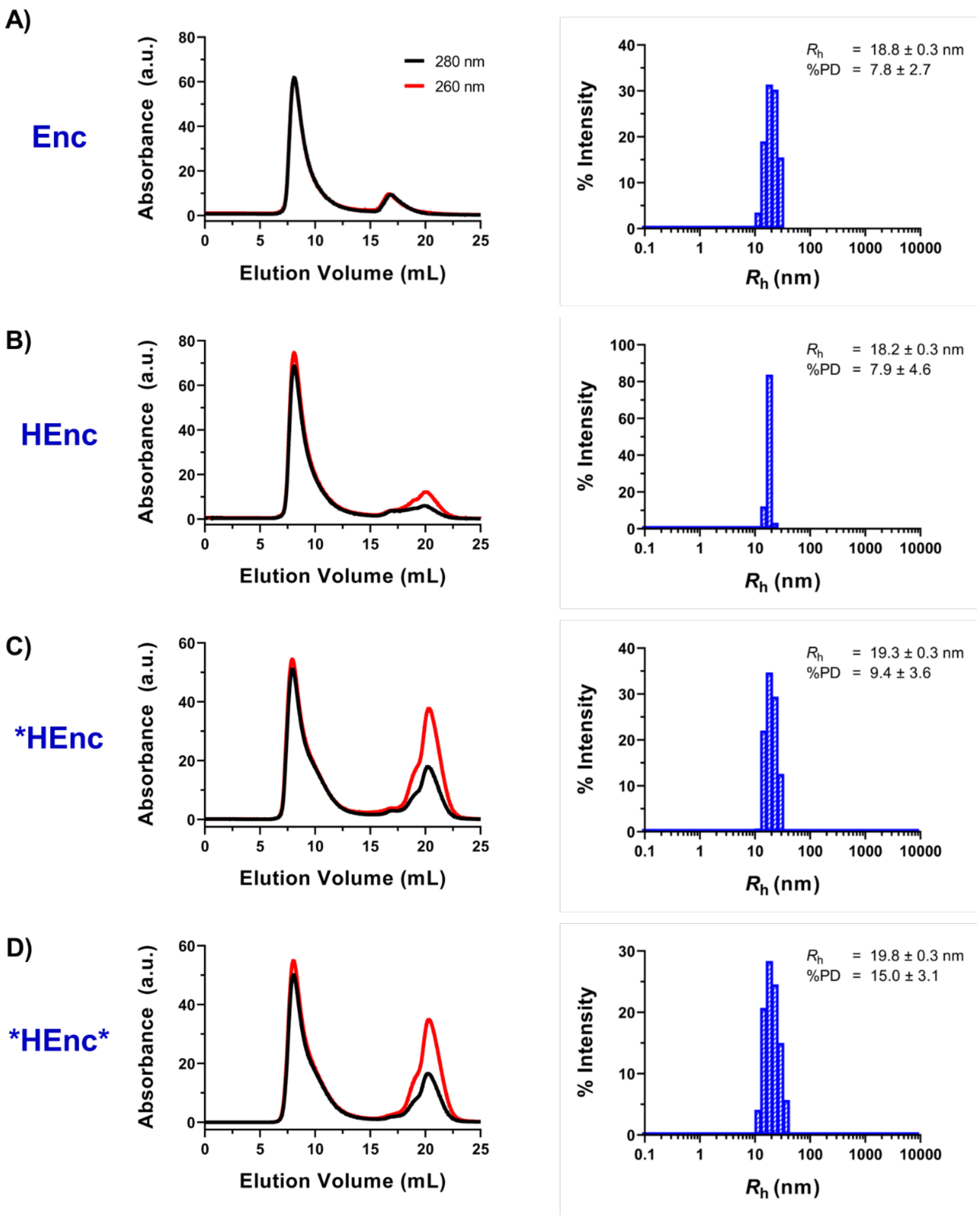

**Figure S2. Characterization of encapsulin variants**. Representative SEC, and DLS data for the A) Enc, B) HEnc, C) *HEnc and D) *HEnc*particles.

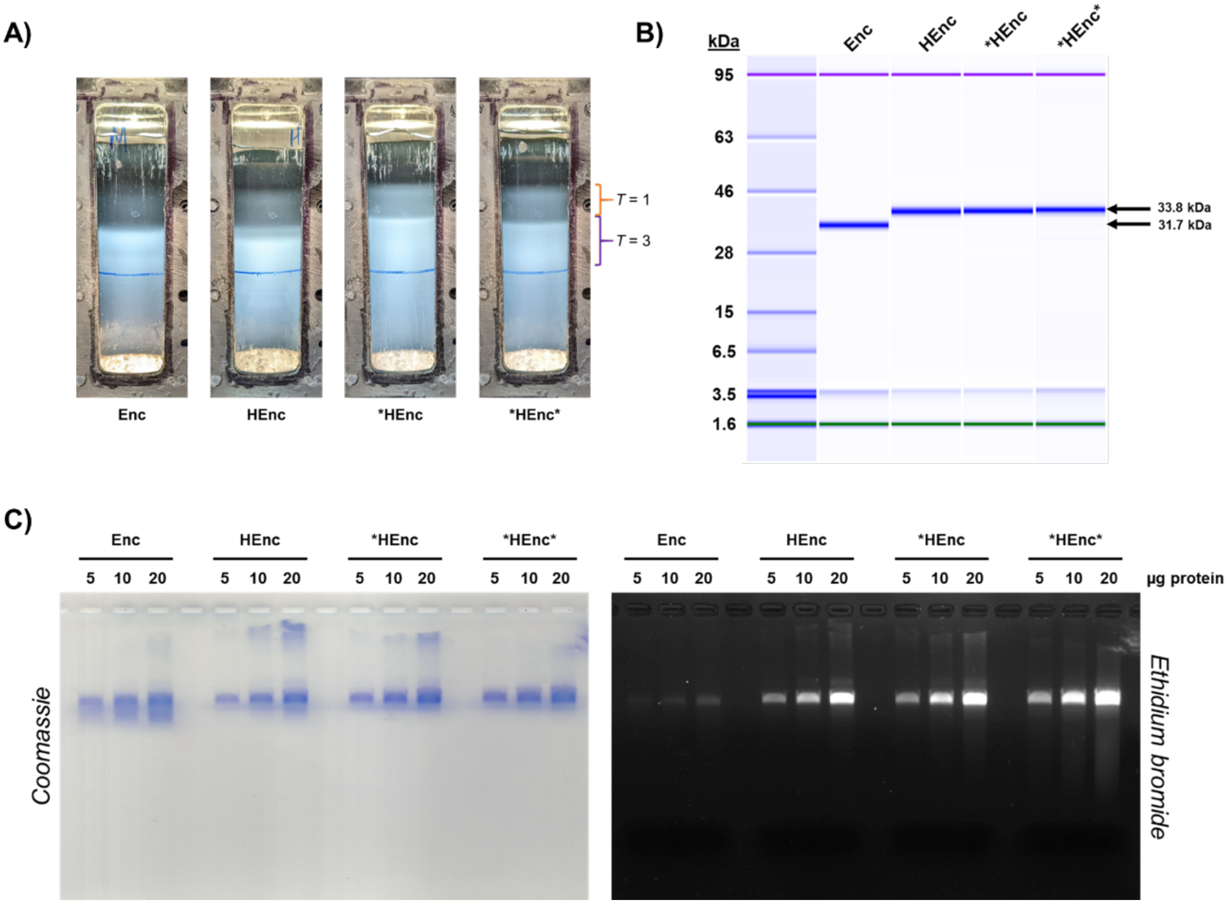

**Figure S3. Characterization of unmodified and RNA-containing variant encapsulins**. A) Sucrose density gradients depicting the $T = 3$ (purple bracket) and $T = 1$ (orange bracket) migration depths after 4 hour centrifugation at 28,000 rpms in a SW-32 rotor. B) Denaturing Bioanalyzer gel electrophoresis assessment of encapsulin CPs. Addition of the N-terminal hexahistidine moiety (full sequence = MGSSHHHHHHSSGLVPRGSH) results in an upward gel shift for the variant encapsulins relative to the unmodified parental CP. C) Native agarose gel electrophoresis assessment of purified encapsulins depicting increased RNA encapsulation for variant PNPs.

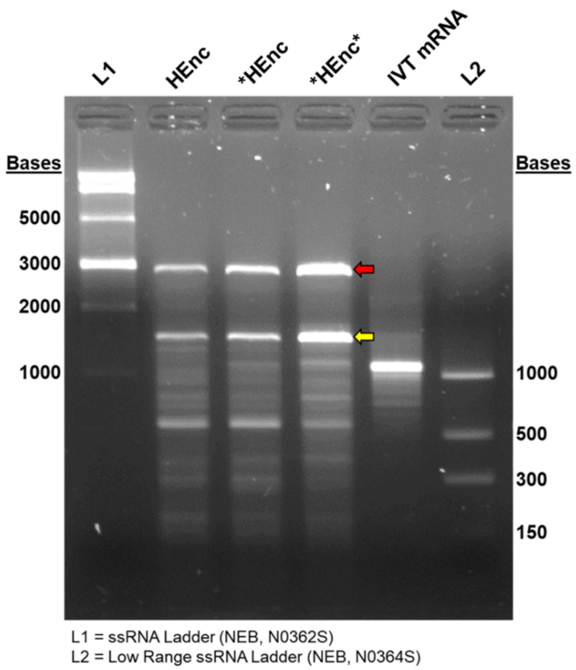

**Figure S4. Denaturing agarose gel electrophoresis assessment of RNAs extracted from encapsulin variants**. *E. coli* 23S and 16S ribosomal RNAs are denoted by the red and yellow arrows, respectively.

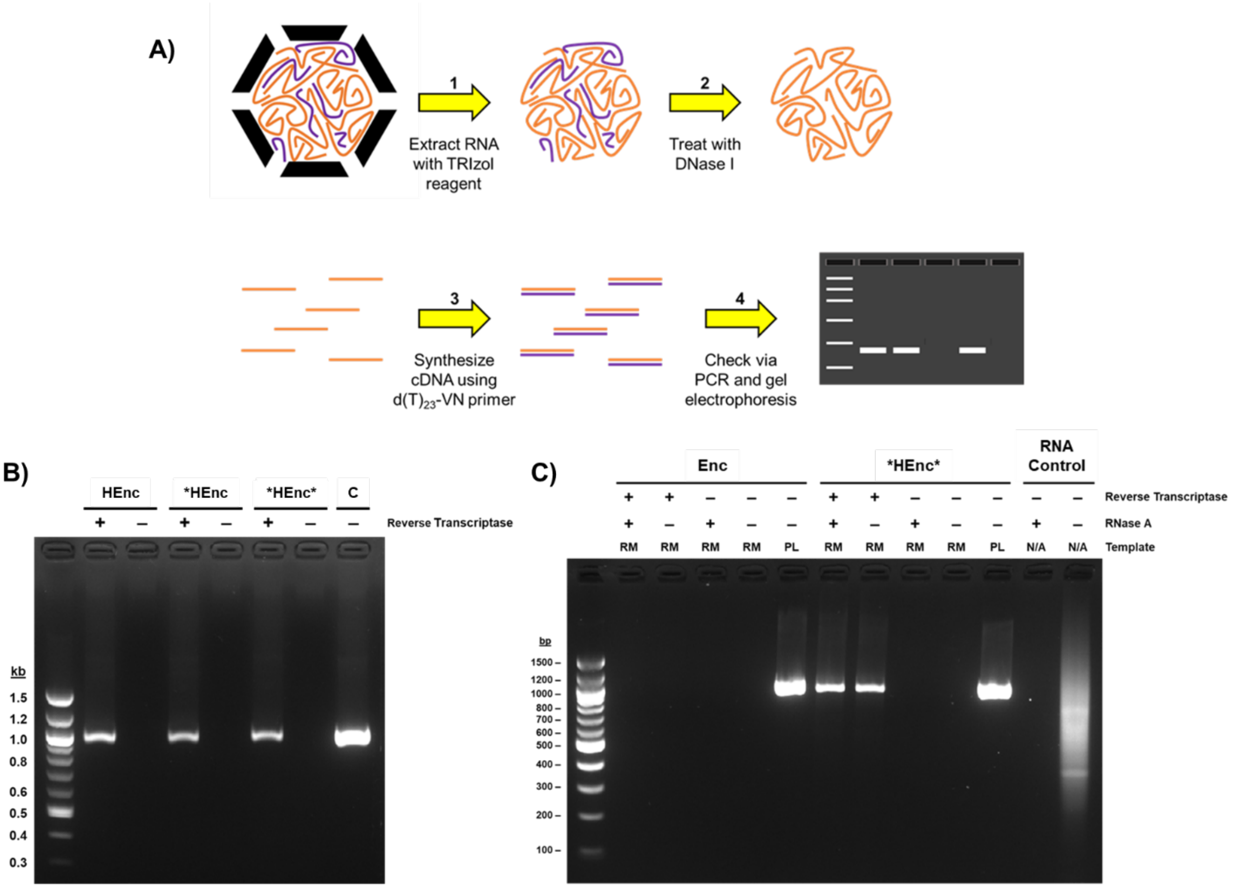

**Figure S5. Confirmation of successful encapsulin mRNA capture within expressed nanoparticles**. A) Overall workflow of RNA extraction from purified nanoparticles, cDNA synthesis, and PCR assessment of nascent cDNAs. B) Agarose gel electrophoresis assessment of PCR DNA generated from cDNA templates using the primers designed to bind to the 5' end of the Enc gene and the plasmid's T7 terminator (expected amplicon = 963 bp). Extracted RNA samples not treated with reverse transcriptase yielded no visible amplicons in the subsequent PCR. C = 10 ng plasmid DNA used to generate a comparative PCR product. C) Control reactions verifying that RNAs isolated from Enc nanoparticles yielded no PCR amplicons following cDNA synthesis, indicating that the unmodified encapsulins package little to none of their parental mRNA. As an additional control, RNAs extracted from both Enc and *HEnc* (as a representative for all encapsulin variants) were pre-treated with RNase A after cDNA synthesis and prior to PCR assessment to verify that the observed amplicons are generated due to primers binding to the cDNAs only and not the extracted RNAs. RM = 2 µL cDNA synthesis reaction mix used as template for PCR; PL = 10 ng plasmid DNA used as template for PCR; RNA Control = 1 µg of RNA extracted from purified Qβ virus-like particles used to verify the enzymatic activity of the RNase A enzyme stock.

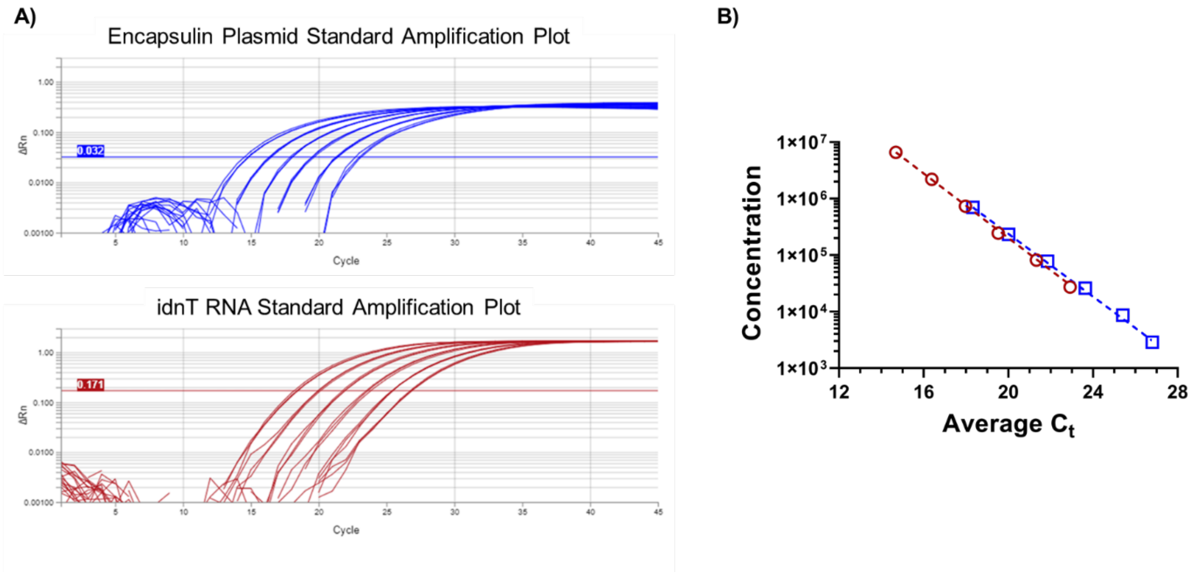

**Figure S6. Example standard curves used for RT-qPCR analyses of PNP-extracted RNAs.** A) Representative amplification plots generated from *E. coli*-extracted plasmids and RNAs with primers targeting the encapsulin protomer gene (blue), and the genomic idnT mRNA (red). B) Standard curves generated from the RT-qPCR amplification plots with affiliated semi-log linear regression fits (dotted lines). The $R^2$ values were consistently > 0.999 for all standard curve regression fits.

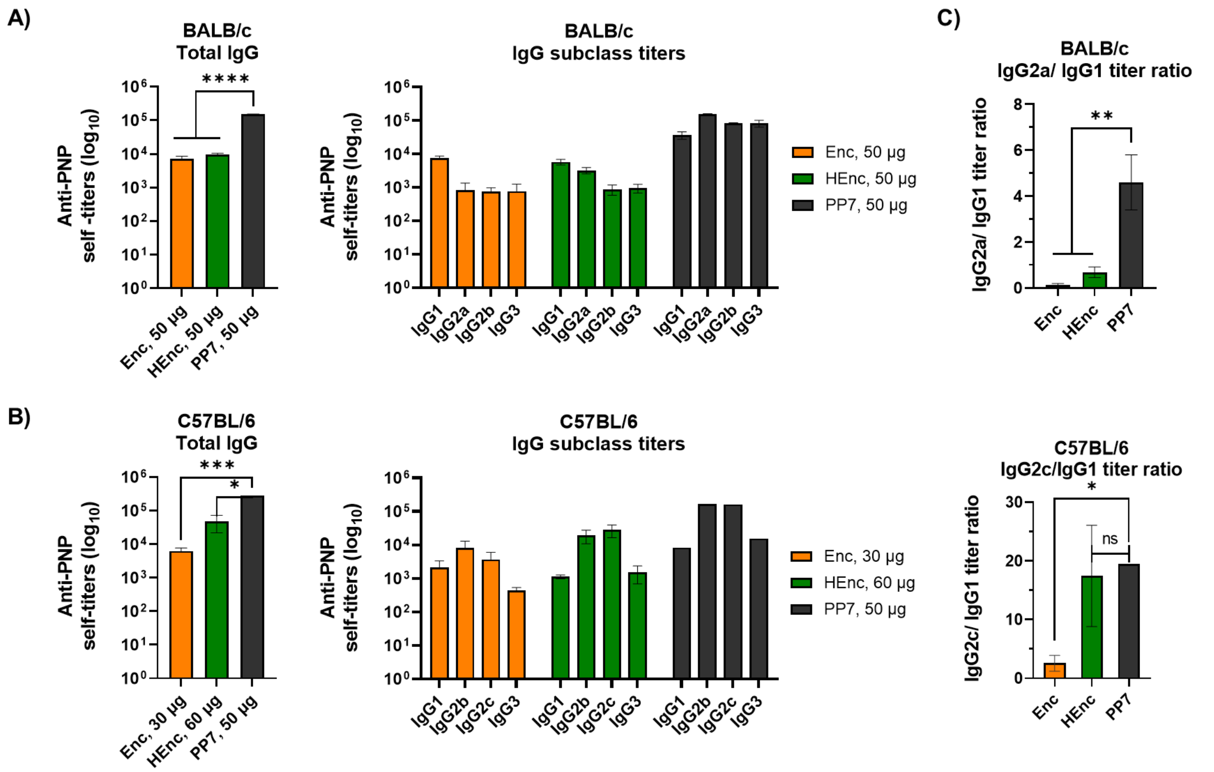

**Figure S7. Anti-scaffold IgG subclasses raised at week 8** in inbred A) BALB/c and B) C57BL/6 mice. C) IgG subclass ratios G2a/G1 for BALB/c and G2c/G1 for C57bl/6. All quantitative data represents shows mean ± SEM and significance analyzed as *=p<0.05; **= p<0.005; ***= p<0.0005; ****= p<0.00005; by two tailed t-test.

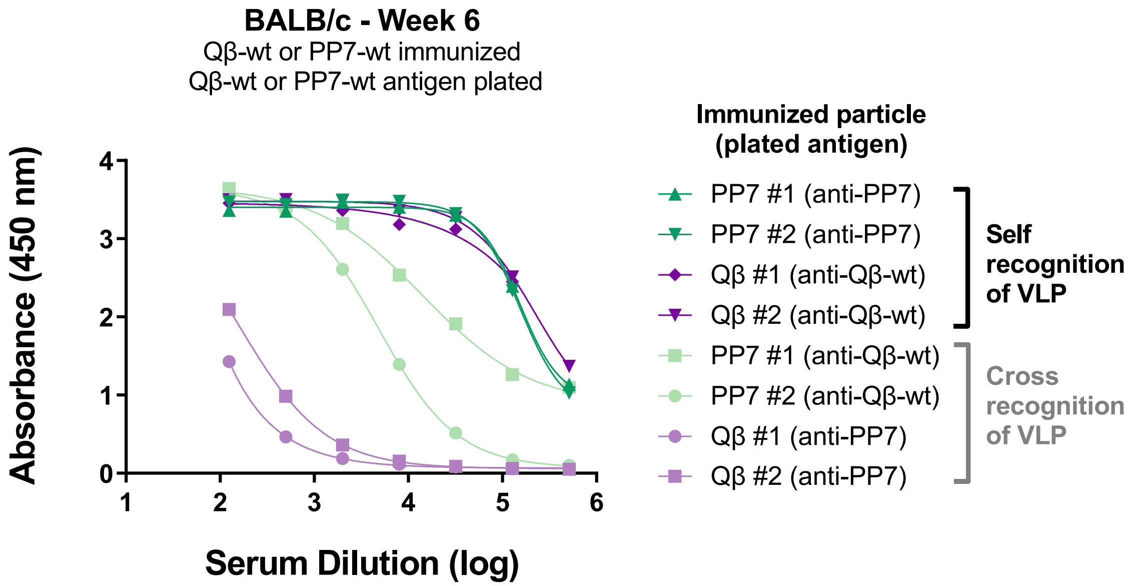

**Figure S8. Cross reactivity of Qβ- and PP7-raised serum antibodies in BALB/c mice**.

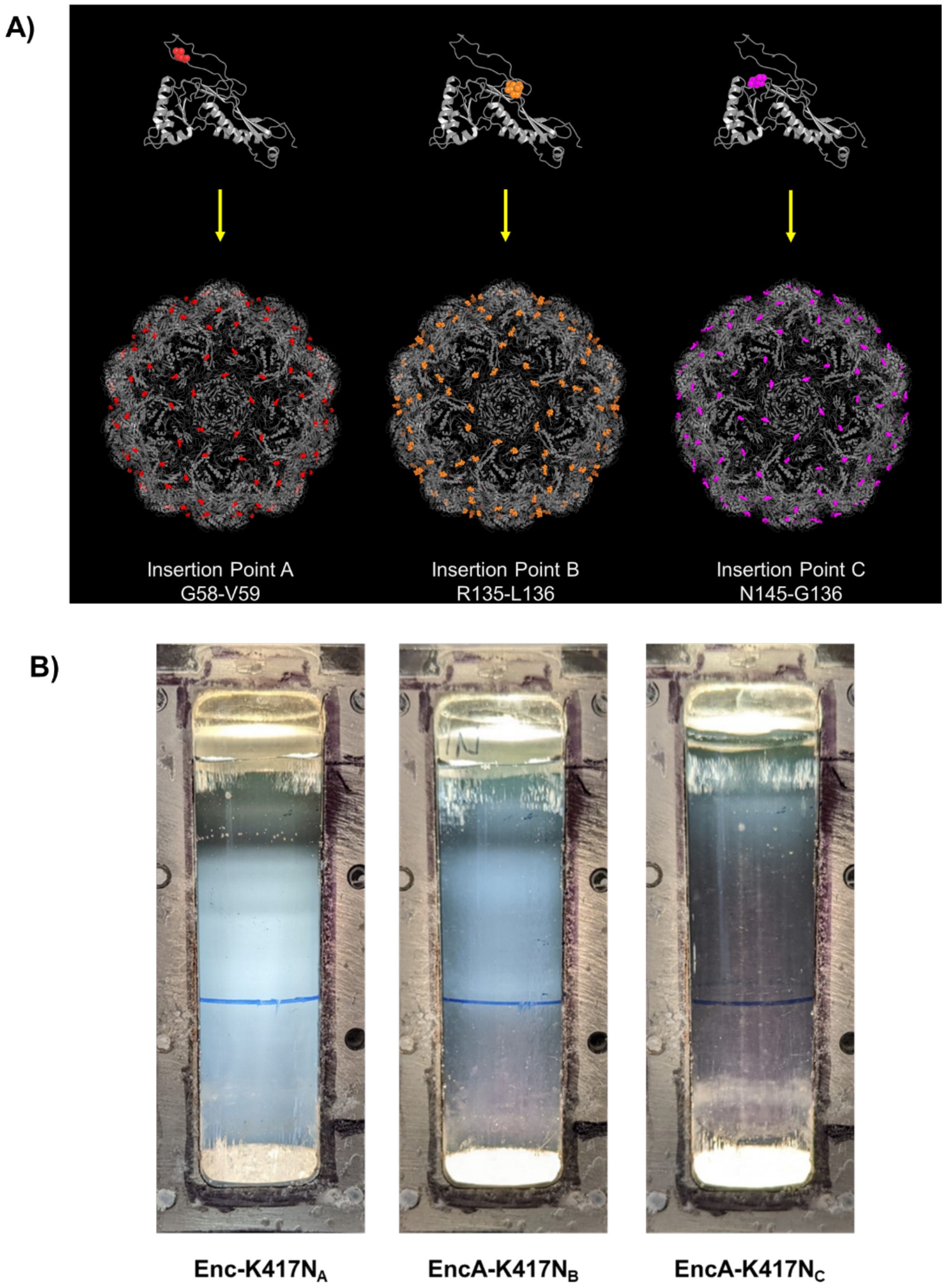

**Figure S9. A)** ***M. xanthus*** **encapsulin CP backbone positions** that served as insertion points for the 20mer K417N peptide (PDB: 7S20). The insertion points are highlighted in the structures above by depicting the residues flanking each insertion point as colored spheres (e.g., residues G58 and V59 depicted as red spheres for insertion point A. The resulting sucrose density gradients produced during the purification of each loop-insertion variant are presented in B).

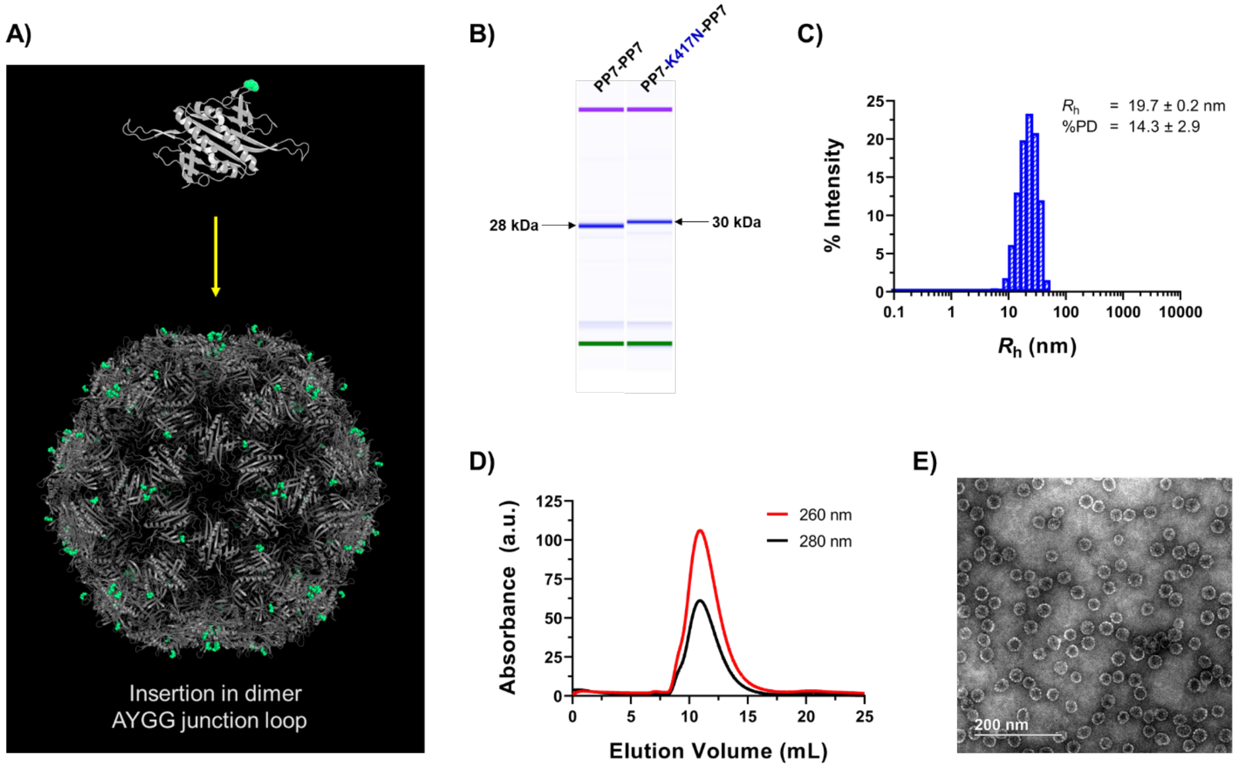

**Figure S10. Characterization of PP7-K417N-PP7 PNPs**. A) Structural representation of the peptide insertion position in the PP7-PP7 scaffold with the two glycine residues of the AYGG junction loop depicted as green spheres. B) Bioanalyzer denaturing polyacrylamide electrophoresis of PP7-PP7 and PP7-K417N-PP7 particles. DLS and SEC plots of purified PP7-K417N-PP7 PNPs are depicted in C) and D), respectively, while a representative TEM image is presented in E).

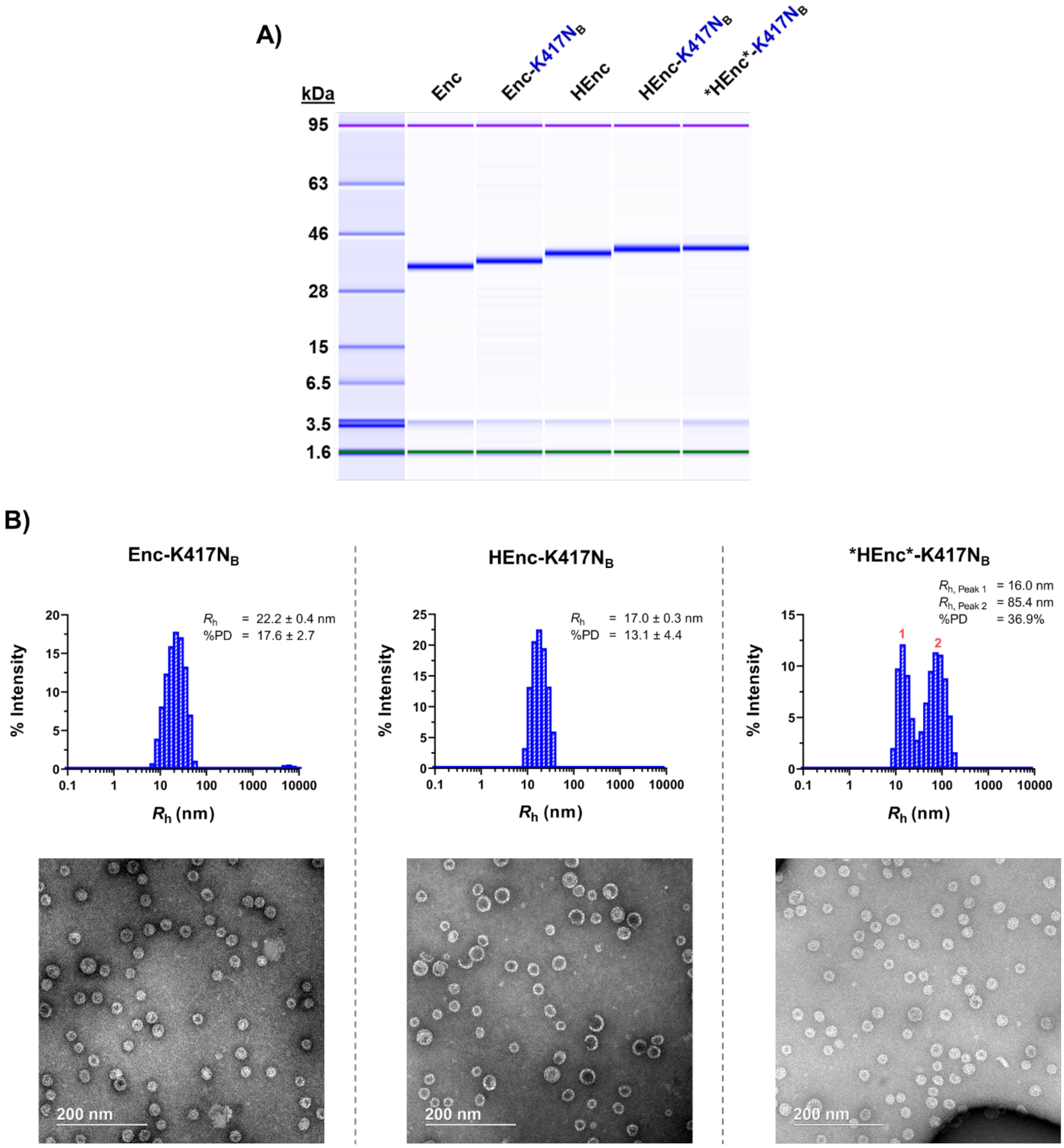

**Figure S11. Characterization of K417N-containing encapsulin variants**. A) Bioanalyzer denaturing electrophoresis of encapsulin variants. B) DLS and TEM characterization of each K417N-containing encapsulin variant. The DLS plot for the *HEnc*-K417N$_B$ sample indicates some possible aggregation of the purified particles in solution, though no apparent particle aggregation was observed via TEM.

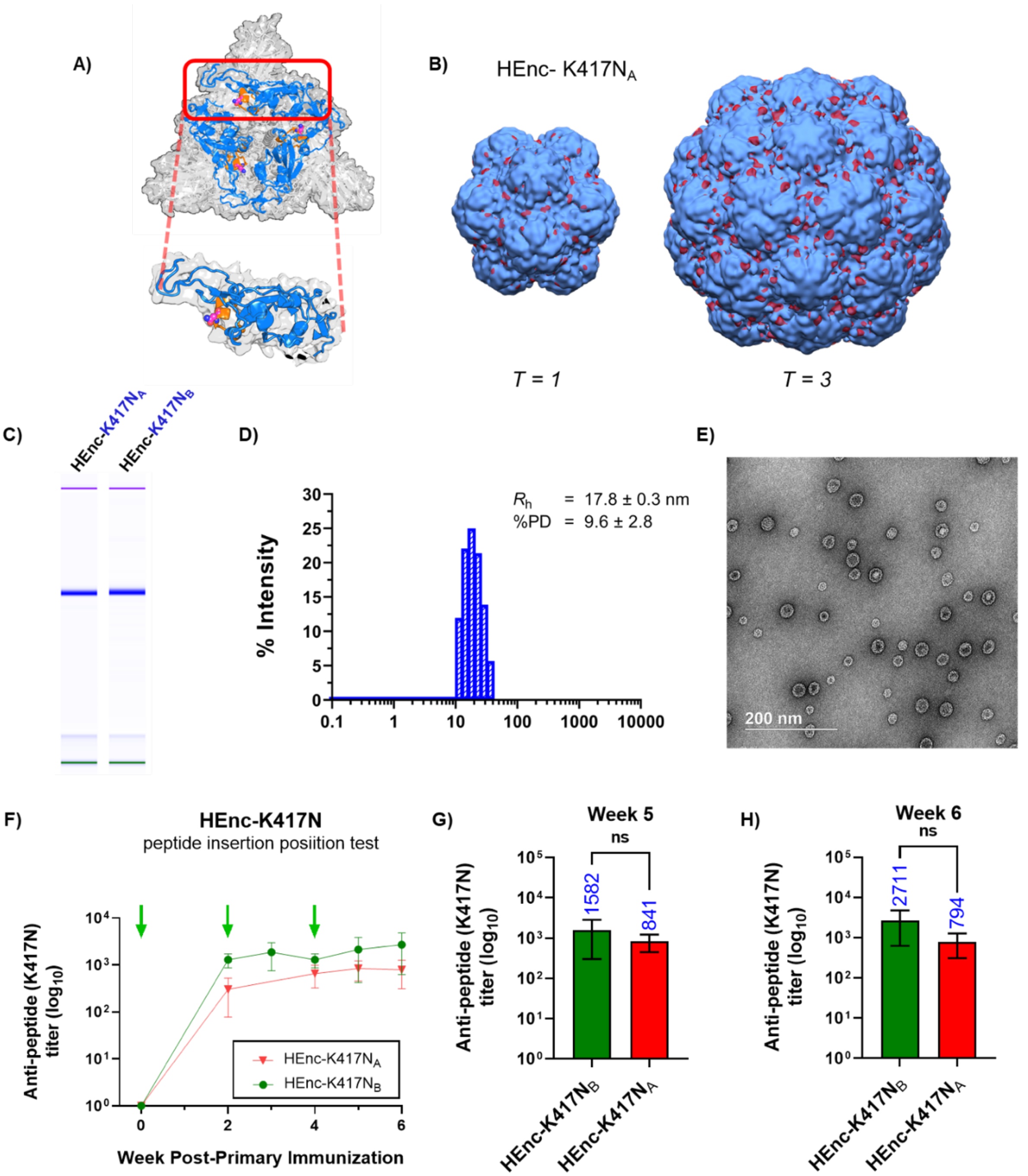

**Figure S12. Comparison of anti-peptide antibody titers generated in BALB/c mice** in response to the insertion position of the K417N peptide in the HEnc coat protein. A) Structure of the trimeric SARS-CoV-2 spike protein with the individual receptor binding domains depicted in blue and the "K417N" peptide sequence depicted in orange. The actual K417N point mutation is depicted with the carbon atoms of the asparagine sidechain represented as magenta spheres. Red box shows an enlarged view of a singular RBD domain. B) Insertion of the 20 amino acid "K417N" peptide into Enc protomers for exterior surface display on PNPs. 10Å resolution cryo-EM electron volume maps depicting the additional electron density present on the surfaces of HEnc-K417NA (red electron clouds) on both $T$ = 3 and $T$ = 1 structures. C) Bioanalyzer comparison of purified HEnc-K417N$_A$ and HEnc-K417N$_B$ PNPs. DLS and TEM characterizations of HEnc-K417N$_A$ PNPs are presented in D) and E), respectively. F) Anti-peptide antibody titers generated by immunization with HEnc-K417N PNPs. More detailed depictions of the week 5 and week 6 titers are detailed in G) and H), respectively.

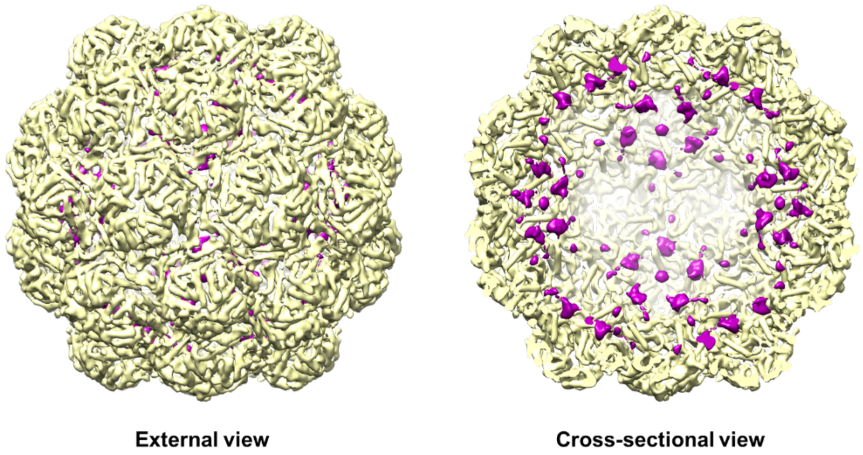

**Figure S13. Cryo-EM volume subtraction map** depicting the additional electron density (purple) present on the luminal walls of HEnc particles as compared to wild type Enc PNPs ($T = 3$ structure shown).

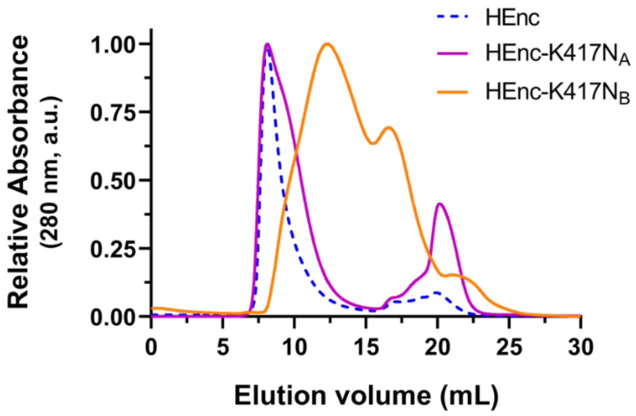

**Figure S14. Comparison of SEC chromatograms originating from HEnc, HEnc-K417NA, and HEnc-K417NB PNP samples**. Both the K417N-containing encapsulins exhibit peak shifts toward later elution times relative to HEnc. For HEnc-K417NA, this effect largely manifests as a broadening of the initial elution peak while for HEnc-K417NB, the initial elution peak nearly disappears entirely, reflective of the drastic reduction in the quantity of the $T = 3$ icosahedral PNPs for this variant.

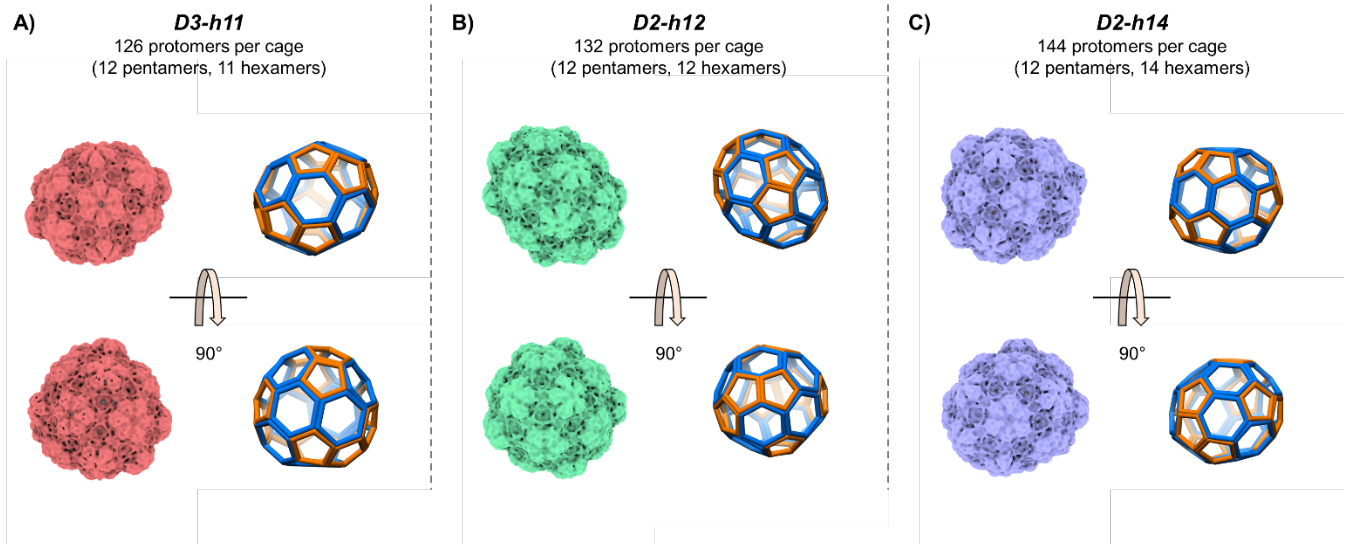

**Figure S15. 7-10 Å volume maps of additional non-icosahedral morphologies observed during cryo-EM analysis of K417N-containing encapsulin variants.** These structures include A) D3-h11, B) D2-h12, and C) D2-h14 assemblies, which were not included in the main text due to their small relatively small numbers within the encapsulin samples. The sparsity of these morphologies likely indicates that they are less stable, and thus less favorable, than the other morphologies that were observed with significantly higher frequencies.

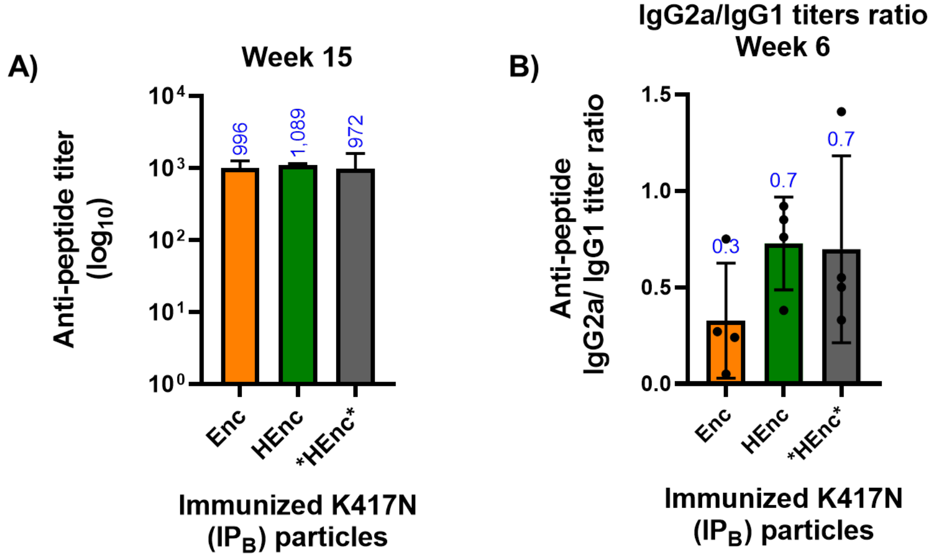

**Figure S16. A) Anti-peptide titers** at week 15 and B) IgG2a vs IgG1 titer ratios at week 6 for all Enc-K417N$_B$ variants.

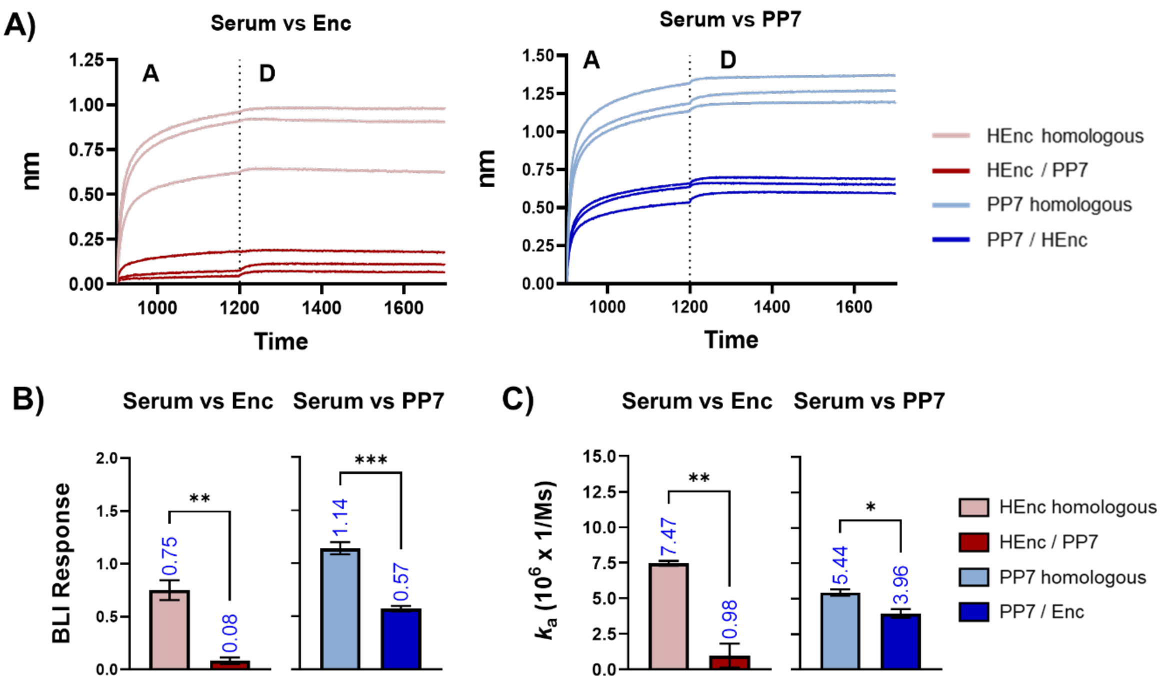

**Figure S17. Biolayer interferometry assays for immunization groups**. A) Serum antibodies at a 1:200 dilution were surface immobilized, then wild type encapsulins (left) or unmodified PP7-PP7 (right) nanoparticles were added in the solution phase at a concentration of 30 µg/mL. Association (A) and dissociation (D) phases are divided by dashed lines in each plot. Antibodies from homologous vaccination groups exhibited greater nanoparticle binding and higher association rates relative to antibodies generated from heterologous vaccinations. Response magnitudes and binding on-rates derived from the BLI kinetic experiments in above panel are depicted in B) and C), respectively. N=3 for each group. All quantitative data shows mean ± SEM and significance analyzed as *=p<0.05, **=p<0.005 or ***=p<0.0005 by two tailed t-test.

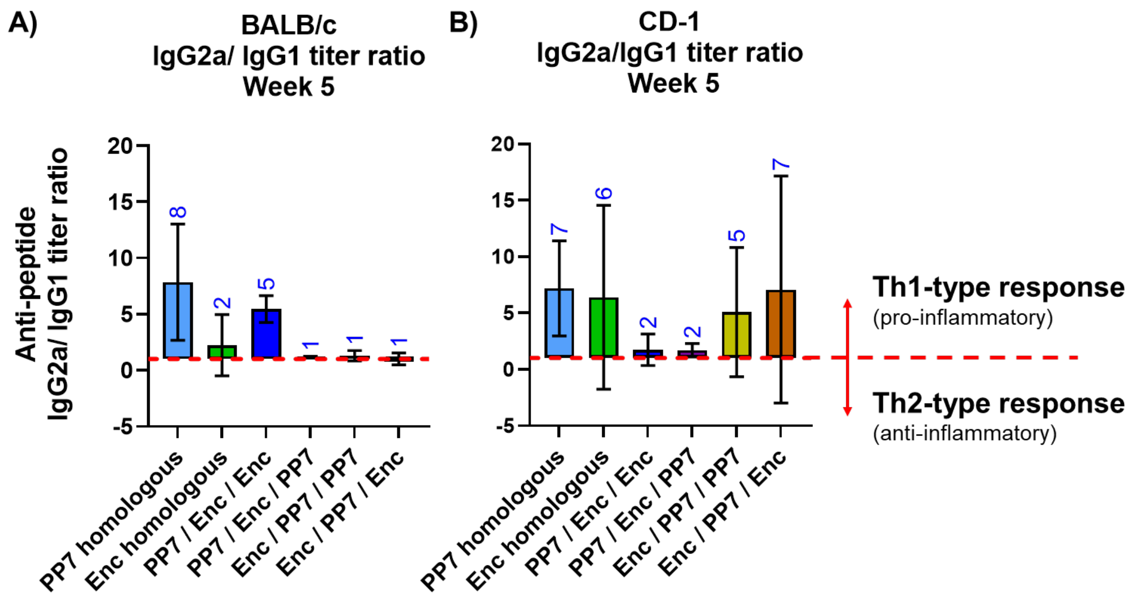

**Figure S18. Ratios of K417N peptide-specific IgG2a vs IgG1 titers** in homologous and heterologous immunized groups for A) inbred BALB/c, and B) outbred CD-1 mice.

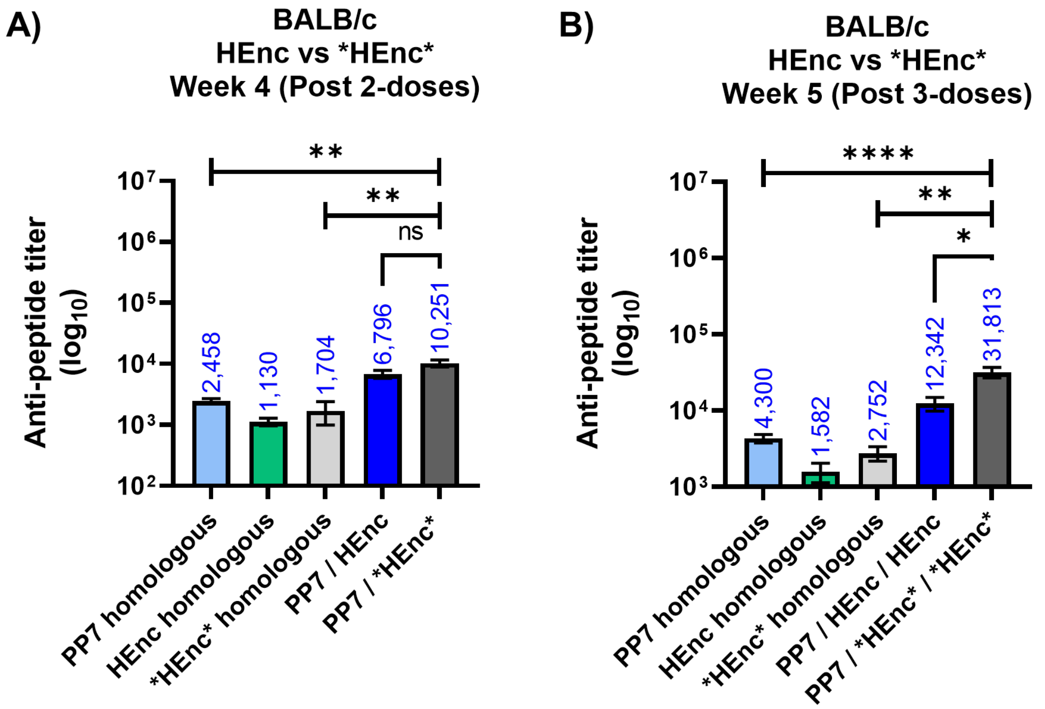

**Figure S19. Improvement of anti-peptide titers** at A) week 4 after 2-doses and B) week 5 after 3-doses by scaffold switching in BALB/c mice. All quantitative data shows mean ± SEM and significance analyzed as *=p<0.05, **=p<0.005 or ****=p<0.00005 by two tailed t-test.

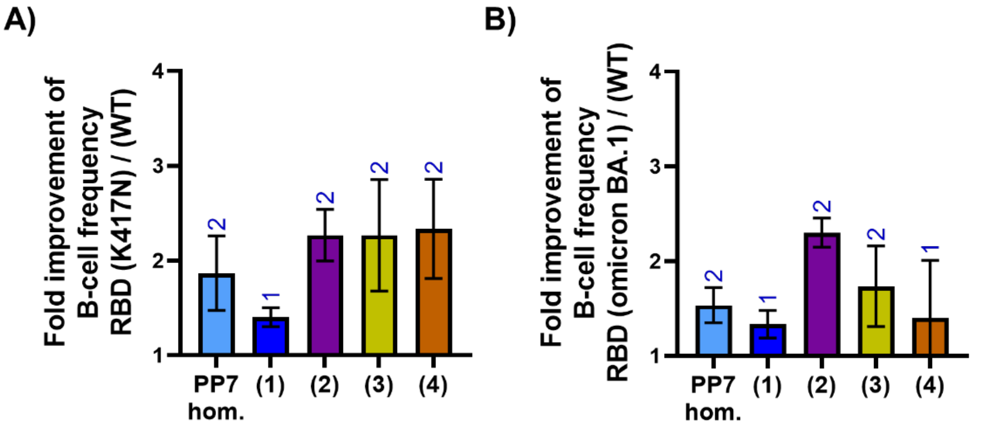

**Figure S20. Fold improvement of frequency of antigen-specific B cell frequencies** of A) RBD(K417N) and B) RBD (Omicron BA.1) over RBD (WT) recognizing cells in splenocytes isolated from immunized CD-1 mice. These plots are derived from the data shown in Figure 7G.

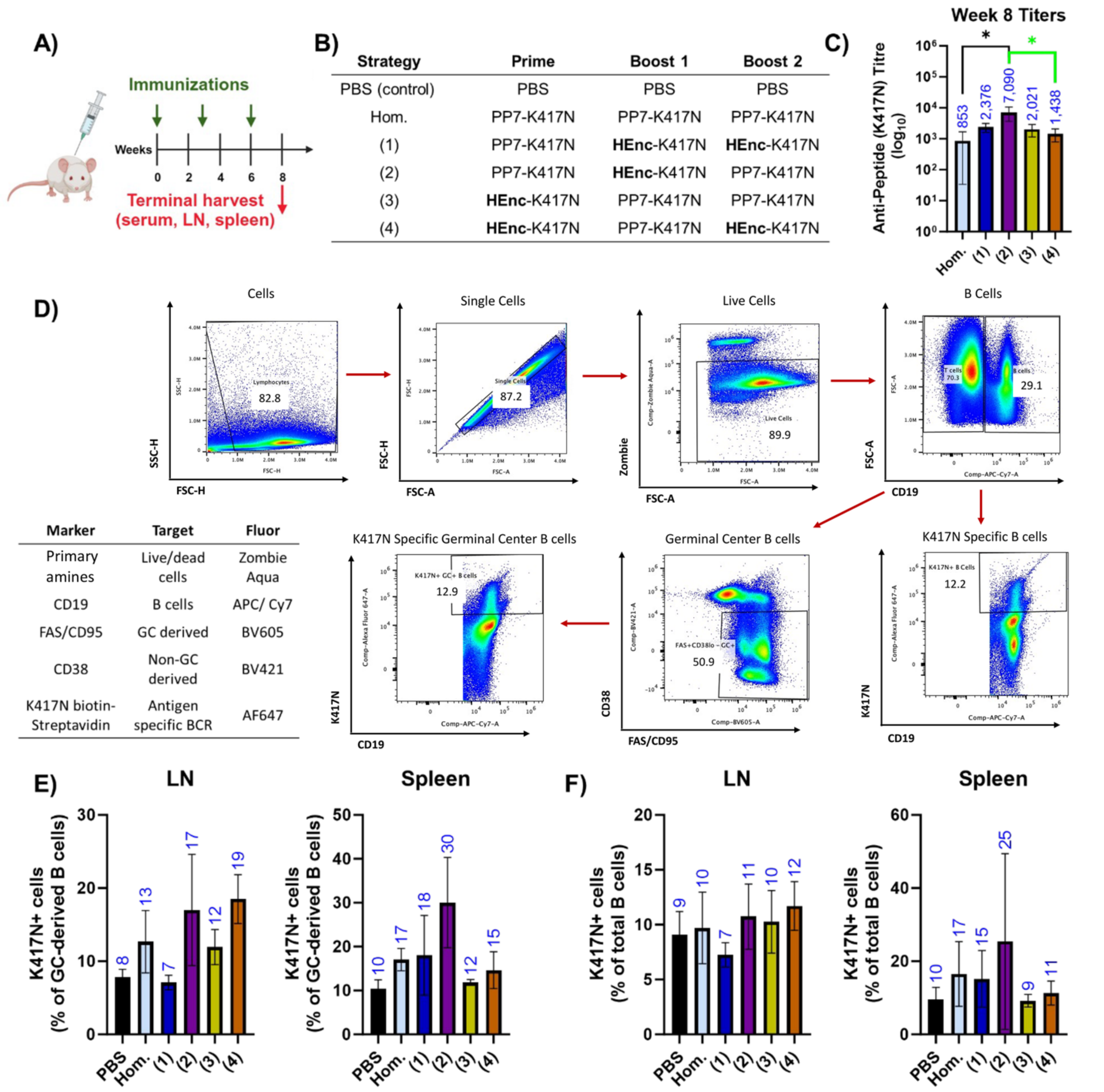

**Figure S21. Assessment of peptide antigen-specific B-cell frequency in lymphoid organs.** A & B) Immunization strategy for heterologous prime-boost in BALB/c mice with three-doses at indicated time point. C) Terminal week 8 anti-K417N peptide serum titers. D) Representative gating strategy for quantification of K417N peptide specific FAS/CD95 + GC-derived and CD19+ total B cells. Inset table indicates fluorophore associated with the targeted cell surface marker used for the assay. Percent K417N-positive population in E) GC-derived B cells and F) total B cells in draining lymph node and spleen. N=4 for each group. All quantitative data shows mean ± SD.

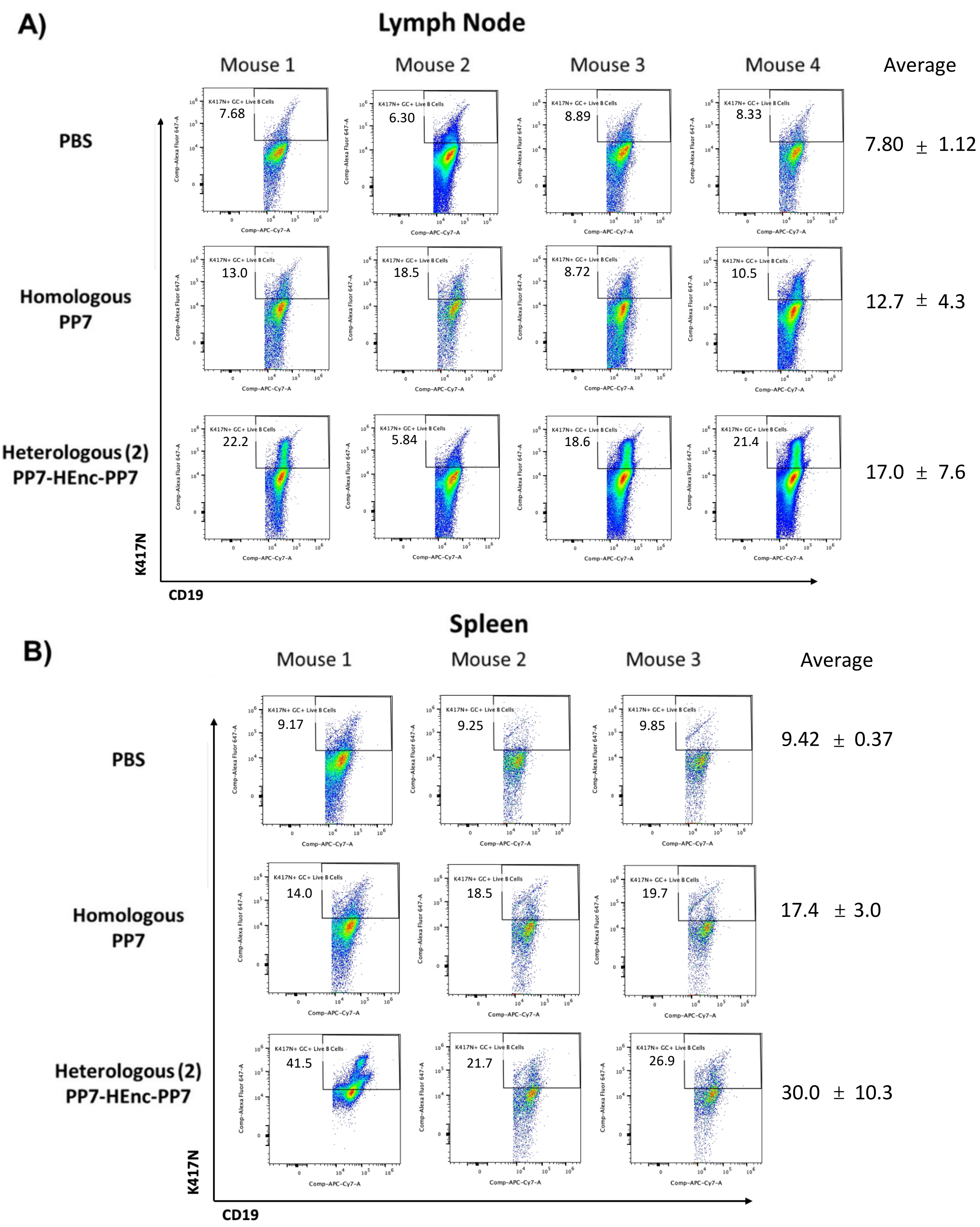

**Figure S22. Representative scatter dot plots** for AF647 labelled K417N positive population in live GC-derived B-cells graphed in panel S21F in organs A) lymph node and B) spleen. Average values (± SD) are shown at the right for each regimen.
